# Longest protein, longest transcript or most expression, for accurate gene reconstruction of transcriptomes?

**DOI:** 10.1101/829184

**Authors:** Donald G. Gilbert

## Abstract

Methods of transcript assembly and reduction filters are compared for recovery of reference gene sets of human, pig and plant, including longest coding sequence with EvidentialGene, longest transcript with CD-HIT, and most RNA-seq with TransRate. EvidentialGene methods are the most accurate in recovering reference genes, and maintain accuracy for alternate transcripts and paralogs. In comparison, filtering large over-assemblies by longest RNA measures, and most RNA-seq expression measures, discards a large portion of accurate models, especially alternates and paralogs. Accuracy of protein calculations is compared, with errors found in popular methods, as is accuracy of transcript assemblers. Gene reconstruction accuracy depends upon the underlying measurements, where protein criteria, including homology among species, have the strength of evolutionary biology that other criteria lack. EvidentialGene provides a gene reconstruction algorithm that is consistent with genome biology.

## Introduction

Unreliability in reconstructed gene sets is an ongoing problem, measurable from missing and fragment orthologs (Trachana et al. 2011; Tekaia 2016; Simao et al. 2015; Waterhouse et al. 2018; Deutekom et al. 2019). Reasons for unreliability include errors from sequencing and assembly, mis-modeling of complex genes, and errors propagated from public databases. One approach that reliably improves recovery of accurate genes is the use of several assembly and/or modeling methods, with evidence-based reduction of the resulting over-assemblies (e.g. Zhao et al. 2011, Gilbert 2012, Haznedaroglu et al. 2012, Schulz et al. 2012, Nakasugi et al. 2014, Smith-Unna et al. 2016, Sahraeian et al. 2017, Cabau et al. 2017, Mamrot et al. 2017, Holding et al. 2018). As these reports show, no single algorithm or tool among many gene modelers and assemblers will produce the most accurate gene set from available evidence. A combination of methods, with varying parameters, recovers more complete and accurate gene sets, by measures of accuracy to reference and to related species genes.

These over-assemblies, or gene sets from several modelers, may contain all of the true gene transcript models. Three general methods of filtering such over-assemblies to remove fragments, mistakes, and reduce redundancy are measured here for recovery of human, pig and plant reference genes. These are self-referential methods, measurements on the intrinsic data of transcripts, and are designed for efficient removal of excess models to produce a draft gene set with all of the accurate models. Such draft gene sets need further, external evidence and expertise to separate remaining error from accurate models.

Three reduction filters compared here are

a. “longest RNA transcript” with global alignment clustering, via CD-HIT (Li & Godzik 2006),
b. “longest coding sequence and alternates” with local alignment, via EvidentialGene (Gilbert 2013),
c. “most RNA-seq” with reads mapped back to transcripts, via TransRate (Smith-Unna et al. 2016).

Filters a, b refer only to the transcript self-sequences, and can be used with any gene sequence collection. Filter c depends on the RNA-seq fragment sequences of those transcripts, genes not in the RNA-seq sample will be filtered out. The three filter tools examined share with peer methods aspects that are common to the type, and specific aspects. Those common and specific aspects are discussed.

Some results of this comparison are obvious: longest transcript filter has longer transcripts, longest CDS filter recovers longer proteins, and most RNA-seq filter yields greater expression measures, compared to the others. The underlying question is which approach returns the most accurate gene information, consistent with efficient reduction to levels at which external evidence can be applied? Where results of these reduction filters differ, the one with greater biological information and phylogenetic validation, is presumed to be of more interest and utility to biologists.

Proteins are evolutionarily conserved, functionally understandable biological information. The biological meaning of coding genes is in their coding sequence, so that discrepancies in CDS versus RNA quality measures favor the CDS measure. RNA-seq expression measures have technical imprecisions, with less direct biological meaning when these qualities deviate from coding sequence quality. The correspondence of protein-related quality measures, including protein size and homology, to biological protein recovery, via proteomics experiments, is known to be well above the correspondence with of expression quality measures (Tress et al. 2017).

This report details use of these three filters to select accurate and complete gene sets from supersets of gene models that contain many accurate genes, plus redundant and less accurate models. Important as well, accurate coding sequence translation is discussed, and the value of several self-referential quality measures for accurate gene set reconstruction. Not considered here are chromosomal evidence, details of homology and external evidence, nor methods of non-coding gene validation. Those are important for accurate gene set reconstruction, and can be applied to the limited-palette results of self-referential draft gene sets. Self-referential gene set reconstruction, when done properly, is an efficient, data-intensive, first step in producing the most accurate animal and plant gene sets.

## Materials and Methods

Materials for this gene reconstruction are RNA-seq published in NCBI SRA database, reference gene sets of human, pig, and arabidopsis, conserved single copy gene proteins, and several software methods, listed in section “Data and Software Citations”.

### M: Reference gene sets

Reference human gene set is from NCBI RefSeq of 2018, with 113,607 mRNA transcripts, containing 81,297 unique coding sequences. Reference plant gene set is for *Arabidopsis* from Araport 2016, with 48,359 mRNA containing 40,880 unique CDS. Reference pig gene set is from NCBI RefSeq of 2018, with 63,593 mRNA containing 47,442 unique CDS. The unique, non-redundant coding sequences are primary reference data used here.

### M: RNA-seq samples

The sampled RNA are paired-end reads, or 100 - 125 bp, from recent model Illumina sequencers. Projects sampled are *human* (PRJNA30709, 98 M read-pairs of SRR3192658), *pig3c* (PRJEB8784, 382 M pairs of 9 runs), and *weed3rd1* (PRJNA323955, 230 M pairs from SRR3664432, SRR3664433, SRR3664434). *Human* sample is from a female cell-line, *weed3rd1* samples of root tissue, and *pig3c* includes adult female and male tissues of muscle, liver, spleen, heart, lung, and kidney.

### M: Transcript assemblies

Transcripts are produced with several assemblers, and various assembly options. This over-assembly of several million models is then reduced to less redundant sets using three types of transcript quality filters. The gene reconstruction pipeline SRA2Genes is used to produce the over-assembly, as described for pig gene reconstruction (Gilbert 2019). Assemblers are Velvet/Oases (Schulz et al. 2012), idba_tran (Peng et al. 2013), SOAPDenovoTrans (Xie et al. 2013), and Trinity (Grabherr et al. 2011). The major option for these assemblies is k-mer size, the sub-sequence length for placing reads in the assembly graph structure. As observed in this and other RNA assembly studies, most k-mer sizes produce some of the best gene assemblies, due to wide variation in expression levels and other factors (Peng et al. 2013, Bankevich et al. 2012). Strongly expressed, long genes tend to assemble well with large k-mer. Alternate isoforms, which share exons and differ in expression levels, are more accurately distinguished at large k-mer sizes (Peng et al. 2013).

### M: Coding sequences from transcripts

Transcripts are translated to proteins and coding sequences with EvidentialGene methods. Evigene ORF calculation methods are compared with several currently used ORF calculations, to ensure they are accurate. Results indicate the methods used are as or more accurate for reference recovery than other transcript ORF calculations in common use.

The ORF calculators compared are NCBI standard ORF finder (Thibaud-Nissen et al. 2013), Evigene cdna_bestorf in two modes, GeneMark transcript protein predictor (2 modes, Tang et al. 2015), and TransDecoder (2 modes). Note that cdna_bestorf and TransDecoder both derive from ORF source code of PASA (Brian Haas et al. 2003), and both have substantial changes for use with assembled transcripts. Comparison of calculated proteins with reference proteins is made two ways: (a) measure and count identical proteins in the two sets, with the reliable and efficient fastanrdb; (b) measure and tabulate complete and subsequence identities, with perl protein string match code.

These command lines are used for calculations, with same input of reference transcripts: **ncbi_stdorf**, “tbl2asn -kc -Mt -Vb -i human18nc.mrna”, where -kc = annotate longest orf, -Mt = for TSA, -Vb genbank flatfile output; **evg_stdorf**, “cdna_bestorf.pl --fullorf --noutrorf --cdna human18nc.mrna”, where --fullorf calls complete protein even when partial is longer, --noutrorf cancels tests for secondary ORFs in UTR left by longest ORF; **evg_bestorf**, “cdna_bestorf.pl --cdna human18nc.mrna”, with default options of “--bestorf --utrorf “; **gmark_gmp**, “genemark_suite/gmsn.pl – euk --faa human18nc.mrna”; **gmark_gmst**, “gmst.pl --faa human18nc.mrna”; **trandec_long**, “Trans-Decoder.LongOrfs -m 30 -t human18nc.mrna”, where -m 30 sets minimum AA to be same as evg, ncbi methods; **trandec_pred**, “TransDecoder.Predict -t human18nc.mrna”, reusing outputs of trandec_long.

### M: Reference alignment

The coding-sequences of transcripts are matched by alignment to reference coding sequences, tabulating all that match at >= 90% identity and within 90% to 110% of reference sizes, as the criteria for accurate models. Tightening these criteria reduces the total number of measured accurate models from the over-assemblies, but it doesn’t change the relative ranking of filter methods. For the complete over-assembly sets, these criteria yield an accurate subset of 32236 models for human, 35748 models for plant, and 33385 models for pig, against which the several filters and individual assemblies are measured for accuracy. In addition, conserved unique proteins are measured in result transcript sets, using OrthoDB (v9, Waterhouse et al. 2013) vertebrate and plant gene subsets, and BUSCO software (Simao et al. 2015), and reported as percent of complete conserved protein set (vertebrate n=2586, plant n=1440).

### M: Alternates and paralogs

Reference transcripts are classified into these gene groups by high identity local alignment of coding sequence, of exon-size or better: **alt** gene groups with 2+ transcripts sharing exons at >99% identity have alternates; **par**, or “paralt” gene groups with 2+ transcripts sharing exons at >97% and <=99% identity have paralogs, and may have alternates; **uni** gene groups have a single transcript sharing no exons at high identity with others. These classes are then used to measure recovery per class, and count, by transcript assemblies.

### M: Definition and use of classify, cluster, and measurement

*Measurements* of transcripts include per-transcript and summary qualities, such as sizes, expression amounts, alignment length and identity, and other items.

*Clustering* brings together transcripts into groups based on one or more measurements (eg sequence-aligned, or multi-variate clusters), but doesn’t distinguish among transcripts in a cluster. E.g. all alternate transcripts of a gene locus share identical exon sequence.

*Classification* separates transcripts into categories, often binary (good/bad, with/without), based on any set of quantitative or discrete measures. For example, transcripts with complete versus partial proteins, with or without homology. Some classifiers may be absolute criterion (e.g. keep only complete proteins), but these have true exceptions often enough that accurate gene classification requires a decision tree or rule hierarchy based on a combination of measures (e.g. keep only complete proteins unless cds is unique and longer than 140aa). Examples of such in genome biology include development of rule-based systems to automatically annotate the unreviewed protein sequences of UniProt with a high degree of accuracy (UniProt Consortium, 2017), and Gene Ontology classifications of gene function.

### M: Redundancy reduction filters

CD-HIT-EST uses clustering by sequence alignment, then classifies the longest per cluster as representative. TransRate is a multivariate classifier, it measures qualities of reads mapped back (RMB) to transcripts, then computes a score from a multiple of four RMB measures for each transcript. Evidential-Gene is a rule-based hierarchical classifier, measuring CDS alignment and other transcript qualities, then applying rules to these measures, including a rule for clustering to gene locus. These filters all classify transcripts into useful (unique, validated or most representative) or not-useful (redundant, fragmented, mis-assembled).

“longest RNA transcript”, with CD-HIT-EST for RNA filtering uses the classification of longest transcript in a cluster of globally aligned transcripts with high sequence identity (identity level is an option). It has virtues and flaws of being a very simple filter algorithm. Similar algorithms are BLAST (local alignment versus global alignment) and several variants. Related methods such as Contig Assembly Program (CAP), Oases-Merge, and others will align-cluster transcripts, then possibly merge them into super-transcripts by overlapping sequence.

“most RNA-seq”, with TransRate uses a classification from expressed RNA-seq mapped onto transcripts, as a multiplicative score of four proportional measurements per transcript: span-coverage *sCcov*, read-map per-base accuracy *sCnuc*, read-pair map agreement *sCord*, homogeneity or segmentation in map level *sCseg*. These RNA-seq alignment measures use specific tools that report read alignment qualities. The full transcript assembly set is evaluated for measures, and a score cut-off for good versus bad transcripts is computed. TransRate classifies the input set into *good* and *bad* transcript sequences.

“longest coding sequence and useful alternates”, with EvidentialGene’s tr2aacds tool uses locally aligned coding sequences to identify gene locus groups, all with identical exon-sized sequence (Gilbert 2013). Locus assignment by CDS overlap in self-alignment is similar to, and gives similar results, as locus assignment by alignment to chromosomes. Locus differences are found for high-identity paralogs that have multiply mapped sequences. It then classifies as useful both the longest coding sequence per gene locus group, and non-redundant shorter alternates. Additional transcript quality measurements are used in this classification, including protein complete/partial, assessment of non-coding versus coding qualities, protein size/complexity, replication across assemblers and samples, and other details that inform gene/transcript model quality. This tr2aacds program implements a classifier algorithm to place transcripts into predetermined categories: primary with alternates (main), primary without alternates (noclass), alternates with high and medium alignment to primary (althi1, althi, altmid), and partial (part) as incomplete alternate transcripts. Alternates can have a fragment qualifier (altfrag), or a protein identity qualifier (a2, aa-high-ident). A classification of *okay* or *drop* is assigned from scores of alignment and protein quality (size, completeness), and optionally protein homology, to partition useful and not useful transcripts.

## Results

Detailed results are provided in a persistent public repository at https://scholarworks.iu.edu/, and at http://eugenes.org/EvidentialGene/other/genefilters_compared/. The EvidentialGene software package is available at http://eugenes.org/EvidentialGene/ and at http://sourceforge.net/projects/evidentialgene/.

An over-assembly of 3,464,117 human transcript models is produced with from one paired RNA-seq sample (98 M pairs) and EvidentialGene’s SRA2Genes, using four RNA assemblers. As noted in Methods, 32,236 of these models match human reference transcripts for the accuracy criteria. Many more match at lower accuracy criteria. An over-assembly of 2,343,913 plant transcript models is produced with from three RNA-seq samples, digitally normalized to 65M pairs, with four assemblers. Of these models, 35,748 match plant reference transcripts for the accuracy criteria. An over-assembly of 8,251,720 pig transcript models is produced with as described for *pig3c* in Gilbert (2019), using the same assemblers. Of these models, 33,385 match pig reference transcripts for the accuracy criteria.

### R: Reference and conserved genes recovery

The transcript classifiers are measured on the reference transcripts of human, plant and pig, as presented in Table 1. The longest CDS method of Evigene retains 94%-99% of non-redundant coding transcripts, longest RNA of CD-HIT retains 68%-87%, and most RNA-seq retains 55%-81%, using a corrected TransRate. The published TransRate software silently fails its intended calculations often, returning only 4% recovery of human reference transcripts, 16% of pig, and 40% of plant. For the *de novo* transcript assemblies, longest CDS method recovers 96%-98%, longest RNA recovers 84%-88%, and most RNA-seq recovers 73%-79%, measured as percent of all accurate *de novo* transcripts. For the subset of conserved unique proteins in these assembled transcripts, with BUSCO/OrthoDB methods, the same ranking is found, longest CDS has highest accuracy, and most RNA-seq has lowest.

**Table 1.** Accuracy at reference gene recovery of 3 general classifier methods. Filter methods: long-CDS = evigene tr2aacds, 2019 update, longRNA = cd-hit-est -c 0.95, mostRNAseq = TransRate published TransRate where noted (^a^), otherwise dgg-patched TransRate. Human_ref, plant_ref and pig_ref are results of filters applied to reference gene sets, human_asm, plant_asm, pig_asm are filters applied to *de novo* assemblies of RNA-seq.

| Filter | Longest CDS | Longest RNA | Most RNA-seq |
| --- | --- | --- | --- |
| Overall Accuracy | highest | middle | lowest |
| *Reference recovery* |  |  |  |
| human_ref | 94% | 68% | 4<sup>a</sup>, 43% |
| plant_ref | 99% | 87% | 47<sup>a</sup>, 81% |
| pig_ref | 96% | 73% | 16<sup>a</sup>, 68% |
| *Reference recovery in RNA Assemblies* |  |  |  |
| human_asm | 96% | 84% | 73% |
| plant_asm | 98% | 87% | 78% |
| pig_asm | 98% | 85% | 70% |
| *Conserved unique genes recovery in RNA Assemblies* |  |  |  |
| human_asm | 96% | 94% | 92% |
| plant_asm | 97% | 93% | 90% |

Supplemental Table AC1 has further statistics, including component assemblers: accuracy at reference recovery is 82% velvet/oases, 73% idba, 73% soap, and 64% trinity.

### R: Reduction filters notes

#### EvidentialGene

The results presented for Evigene are from an updated software release (2019.10, v4). The Evidential-Gene method tr2aacds, as of 2018 (version 2, 3), did not reach expected recovery of reference alternates and paralogs, although it was found superior to other filters. Algorithm lossage was examined in detail, and corrections made to be consistent with retaining true reference alts and pars, while maintaining redundancy reduction. See below and Supplemental Document EVUP for details of evigene’s gene transcript quality filter using self-referential methods, prior and updated algorithms, with code and parameter changes explained.

#### CD-HIT-EST

CD-HIT-EST with default options failed, or took excessive compute time, on all large transcript set reductions, for the sequence identity default of -c 0.90. Instead the option -c 0.95 is used here, as commonly reported in various studies, and works reliably for large sets, though at ~10x greater compute cost than tr2aacds.

#### TransRate

TransRate software as published results in very low recovery of reference transcripts, as noted in Table 1. This TransRate release fails in silent and somewhat complex ways that requires expert diagnosis. This happened for this author’s trials on several Linux and MacOS systems, and is related to binary code versions, and missing pipeline checks. One diagnostic of these failures is a transrate assembly score of zero, reported in various publications (e.g. Smith-Unna et al. 2016 Sup. Table 2; Holzer & Marz 2019). No TransRate software updates have been provided since 2016, and it doesn’t readily rebuild from sources. This author has patched source code and updated the salmon component for a version that works without silent failures; see Supplement TRFIX text for details of these code changes. This patched TransRate worked for human and plant samples, but failed on the larger pig over-assembly, due to component software limits. A smaller set of 3.4 million pig transcript models, those with non-redundant coding sequences, is analyzed and reported, but some measures are not comparable with the other two filters (i.e. filtered transcript count).

**Table 2.**
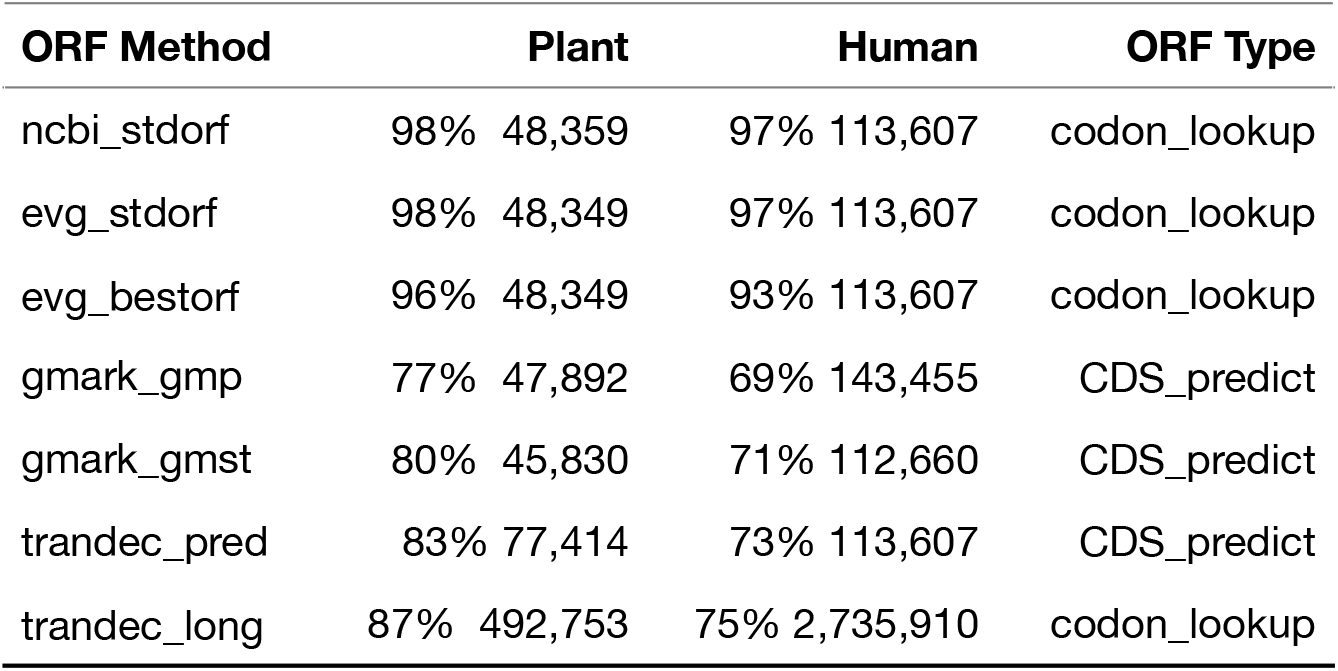
Recovery of reference proteins by ORF computations from human and plant reference mRNA. Percentages that match reference proteins, and counts of ORFs computed from reference transcripts are given. Methods in brief are, ncbi_ and evg_ of NCBI tools and EvidentialGene, using a standard codon lookup table, gmark_ and trandec_pred of GeneMark and TransDecoder, using prediction methods, and trandec_long, from codon lookup. ORF methods, and human and plant *Arabidopsis* reference sequences are detailed in Methods.

At least part of the difficulty with TransRate is measurement of multiply mapped reads, which are valid components in alternate and paralog sequences. A valuable diagnostic of errors in TransRate, or any RNA-seq based measure of transcript quality, is measurement of reference genes distinguished by alternates versus uniques; the transrate score drops for the same transcript when measured in presence of alternates at that gene locus, but not for unique-sequence transcripts. E.g. for *Arabidopsis* reference, the average score for longest transcript of loci with alternates changes from 0.144 (low) to 0.329 when alternates are removed from analysis, compared to constant score of 0.25 for gene loci with one transcript. Three score components drop in presence of alternates: sCnuc, sCcov, sCord, while sCseg rises. The TransRate publication does not report or discuss effects of alternate transcripts, paralogs, or RNA-seq multiple mapping measurements, yet these are critical in distinguishing valid biological transcripts from assembly artifacts.

### R: Coding sequences from transcripts

Computation of proteins and coding-sequence span in transcript assemblies is an essential method for gene informatics, one that is simple at its basics: the use of well-researched codon translation tables. Although this methodology is well-known from early days of bioinformatics, there are modifications, especially related to distinguishing valid proteins from sequence with ambiguous coding potential or short random codes. Current practice in transcriptome informatics includes tools that predict coding potential and coding sequences in various ways. Some of these tools, though popular, return results that are not consistent with reference gene sets. This report compares several currently used tools for proteins, or open reading frames (ORF), calculated from transcripts; these are described in Methods. The main distinguishing aspect is whether these are primarily at codon table lookup method, where ORFs are computed by scanning transcripts in 6 offset frames, collecting codons from start to stop, and determining the longest ORF usually as best choice. This choice usually matches experimental results (Tress et al. 2017), but exceptions occur in biological translation to proteins. The other aspect here involves prediction, using coding potential, sequence signals beyond the codon table, and other prediction methods, attempting to separate likely translations from unlikely ones.

Table 2 summarizes the answer to this basic validation of whether ORF computations reproduce reference proteins from their transcripts. Standard ORF computations that NCBI and others employ, including Evigene, do reproduce reference proteins. The prediction methods do not do so, mis-calculating 17% to 30% of reference proteins. GeneMark (Tang et al. 2015) and TransDecoder are mis-calculating reference proteins for various reasons, in different ways. A large portion of the mis-calculations are as longer, partial proteins than reference. This happens when mRNA contain a 5’ UTR leader that has valid codons, but extends beyond the start codon, often experimentally determined. Biological reasons for such are known (eg. uORFs and oORFs, Johnstone et al. 2016, Juntawong et al. 2014). While longer than true protein is less of an error than no translation or mistaking most of the codons, such too-long errors do affect their accuracy value. Supplemental Document OCC describes computational details for these ORF calculators, and details of how they are mis-calculating reference proteins.

The EvidentialGene method ‘cdna_bestorf.pl’ has options that affect its best-guess when 5’ end contains a longer partial protein: in standard mode, it matches all but a few percent of reference proteins, in its ‘bestorf’ mode it guesses based on how much longer the partial end is, whether to choose a complete or longer partial protein. All these methods miss a few exceptional proteins, including ones that require experimental or expert assessment, or are shorter than minimum sizes of 20aa-100aa (Evigene uses 30aa by default, NCBI 20aa to 30 aa). One outcome of this assessment is the cdna_bestorf default parameter for 5’ partial proteins is reduced, to avoid many of these extension errors.

### R: Alternate and paralog recovery

A standard method of classifying transcripts to gene loci is by mapping, or modeling, onto chromosomes, with the criterion that transcripts at one locus share some coding exon locations. Paralog genes do not share coding exon locations, by common definition, though their exons may be sequence-identical. EvidentialGene instead classifies loci by alignment of coding sequences. This gives nearly the same results as mapping to chromosomes, though high identity paralogs are confused as alternate transcripts, measured at 3%-5% of paralogs for animal and plant reference sets. A smaller 1% portion of alternates at one locus are misclassified as paralog loci. *De novo* gene assembly methods that classify loci have similar discrepancies, as RNA-seq reads are often identical among paralogs. A recent report with methods developed to disentangle artifact from biology finds biological evidence of cross-paralog sharing of coding exons (Dougherty et al. 2018), in exception to the common definition, but not a surprise to this author with experience trying to disentangle tandem duplicate genes in a range of species.

For this report, reference transcripts are classified into gene groups with categories of **alt**, **par** and **uni** as noted in Methods, using local identity of coding sequences. This permits measurement of accuracy in categories, an area that is under-reported for transcript methods, and as this report shows, where they diverge significantly. This computed category agreement with reference-supplied loci of human, pig and plant, is **alt**: 99%, 92% and 98%, **par**: 86%, 91%, 57%, and **uni**: 100% for all three. The lower agreement for paralogs is for those with identical coding exons. For purposes of this report, the computed categories are sufficient, as trends are the same for both categories, and are reasonably interpreted as effects of sequence identity levels.

The measured accuracy at gene category recovery is summarized in Table 3, for human and plant gene groups. For the unique transcripts, Evigene recovers all but 3%, while CD-HIT and TransRate miss up to 20%. For categories with multiple transcripts, Evigene is somewhat reduced, to 90%-94% for two transcripts, and 80%-90% for four alternates or paralogs. The longest RNA filter drops from 70%-80% of one alt/par to 56%-67% for two, down to 34%-48% for four alt/par transcripts. TransRate’s most RNA-Seq retains very few accurate alt/par transcripts, below 10% for four, as it appears to suffer from mis-measurement of multi-mapped RNA-seq that is common to those. Linear regression statistics indicate a significantly higher intercept and slope to the accuracy of Evigene, and higher intercept for CD-HIT over TransRate, for these results (Table 3, Figure 1, statistics in Supplement AAP).

**Figure 1.**
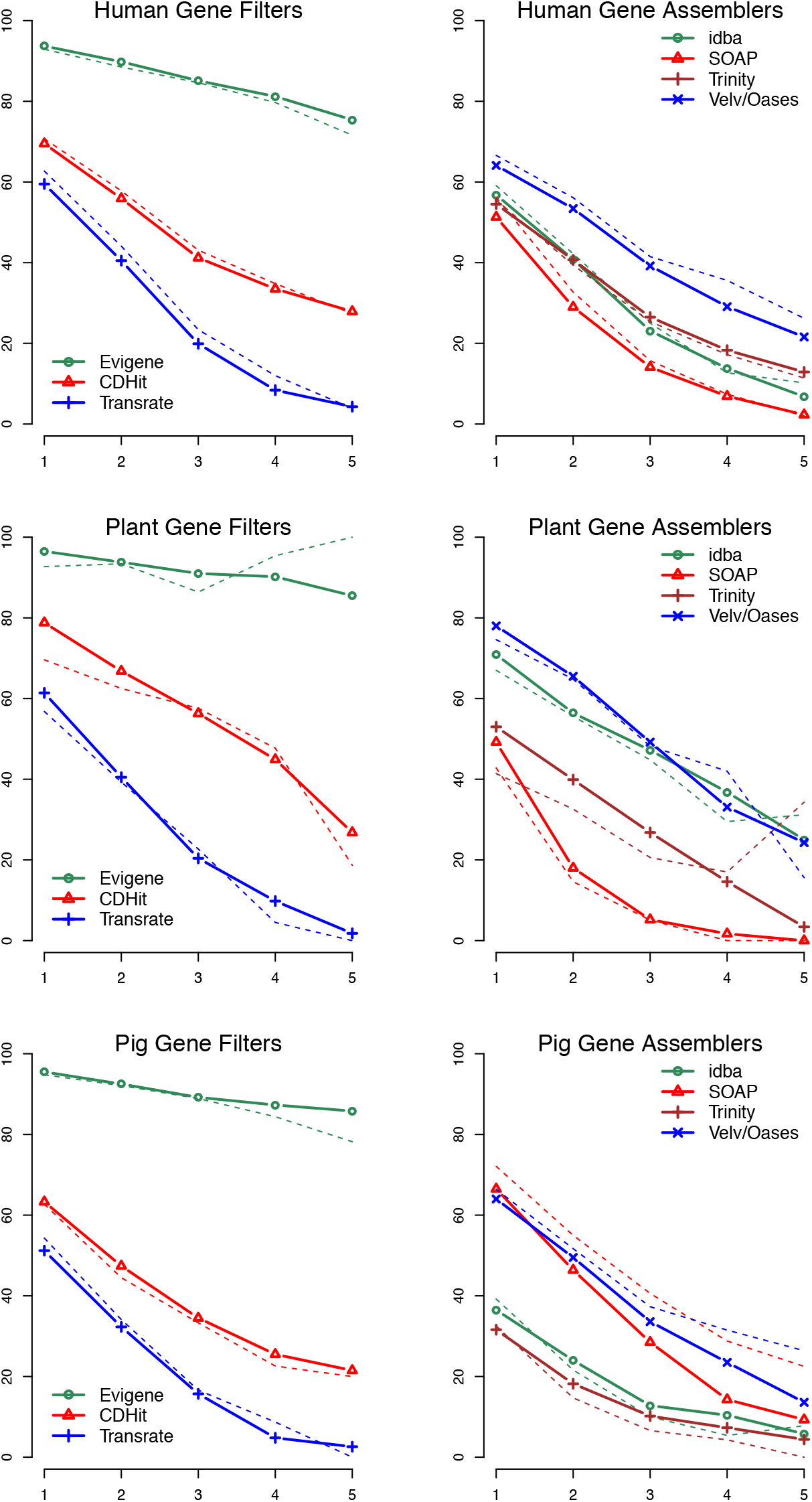
Reference recovery for human, pig and plant gene sets, comparing reduction filters and assembler methods. Y-axis is percent recovery of available reference transcripts, x-axis is number of transcripts accurately recovered in gene families (alternates in solid lines, paralogs in dotted lines). Not shown are unique gene loci (1 transcript per reference family).

**Table 3.**
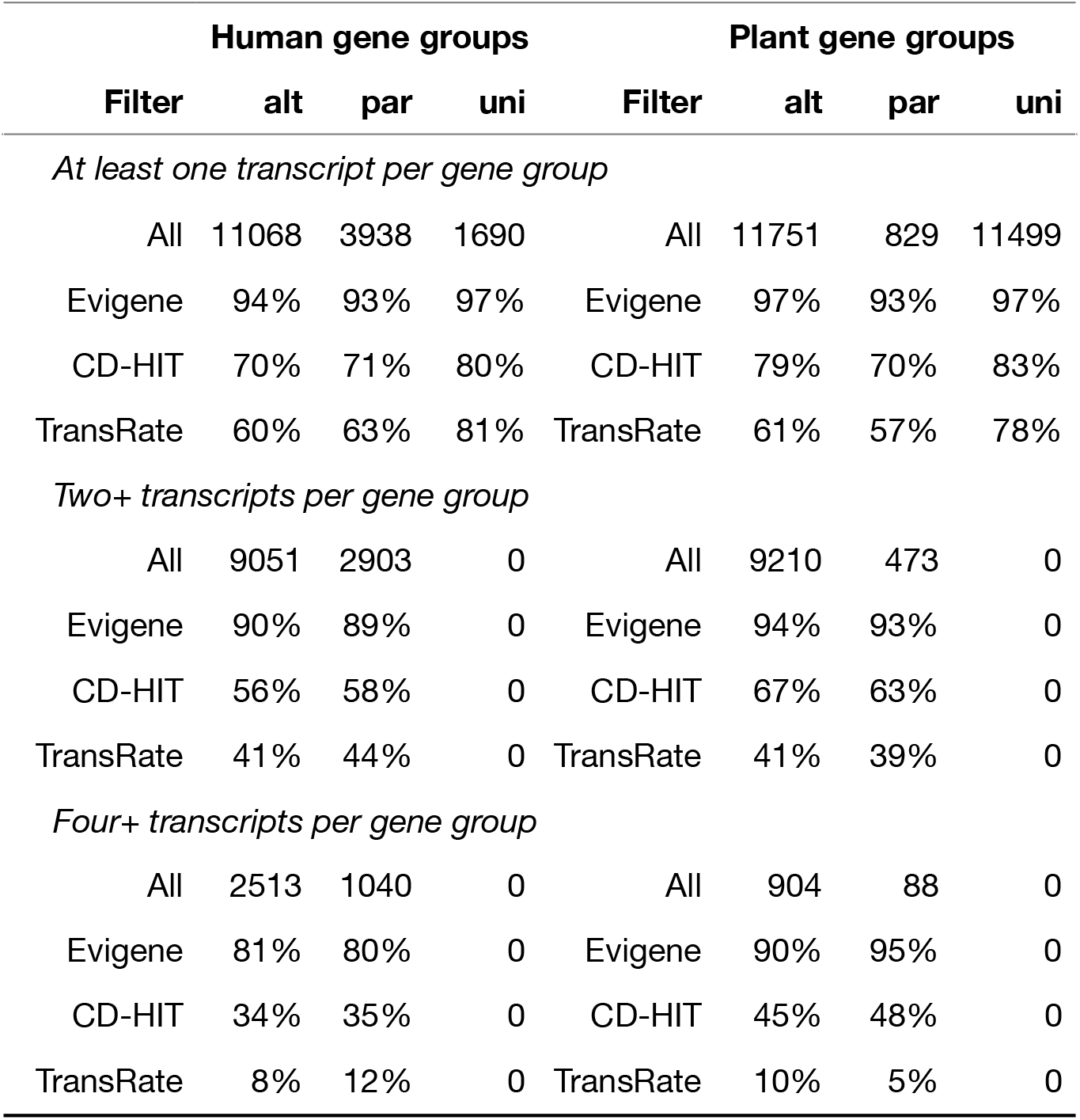
Accuracy at recovery of reference alternate and paralog transcripts, for human and plant *Arabidopsis* reference gene sets. These are recovery in transcript assemblies of the reference transcripts, measured by alignment at criteria of >90% identity and size. Filter types are *All*, counts of recovery in all of the transcript assemblies, at criteria, or percent of all when filtered by Evigene, CD-HIT, and TransRate methods. Group types are **alt** = gene group contains alternate transcripts, **par** = gene group contains paralogs (and alternates possibly), **uni** = gene group contains single, unique reference transcript. See Figure 1 for plots of these and related recovery percent per transcript count/group.

Recovery of alternates and paralogs by reduction filters, as well as individual assemblers, are plotted in Figure 1, as percent recovery of the total assembly set’s accurate transcripts, in relation to the number of reference transcripts in **alt** and **par** categories. Evigene’s CDS filter is the most accurate, and preserves its accuracy at high levels with the additional alternate and paralog transcripts. Notably, none of the 4 individual assemblers preserve the additional accurate alternates and paralogs that are recovered from their combination. This is in part a matter of options used for each assembler, as at least 3 have options that will increase numbers of alternate transcripts they report. All however were run with such options, and report many alternates, but ones that do not individually agree as well with reference transcripts.

The accuracy of Velvet/Oases assemblies is regularly, and significantly, higher than with Trinity and some of the other assemblers, though as found here and elsewhere (Zhao et al. 2011, Haznedaroglu et al. 2012, Schulz et al. 2012, Yang & Smith 2013, Sahraeian et al. 2017), results for assemblies vary with options used, species, data set and measurement qualities. Supplemental Document AAP has counts, percentages of agreement, and statistical analyses of results used in Table 3 and Figure 1.

### R: Other reduction filter qualities

Basic quality statistics of these reduction filters include transcript size (eg. N50), protein size (eg. AA1K), and expression (eg. coverage span, depth or level). All three reduction filters are reducing the over-assemblies of 2 million to 8 million transcript models by about 90%, to a usable draft gene set for further work, as shown in Table 4. None of these have produced a desired “same as known” gene count (see Discussion). The count of filtered transcripts shows that Evigene’s longest CDS approach is reducing redundancy well, while retaining most of the accurate transcripts.

**Table 4.** Other quality statistics of filter-reduced gene assemblies. These quality statistics are detailed in text. The “best” quality values are bolded (lowest for count, highest for others). Filter counts are around 10% of the total over-assemblies of 3,464,117 human transcript models, 2,343,913 plant transcript models, and 7,788,347 pig transcript models.

| **Filter** | **Longest CDS** | **Longest RNA** | **Most RNA-seq** |
| --- | --- | --- | --- |
| *Filtered count* |  |  |  |
| human | **448380** | 526884 | 578882 |
| plant | 297327 | 560938 | **252062** |
| pig | **869646** | 938953 | na |
| *AA1K Protein size (aa)* |  |  |  |
| human | **3527** | 2684 | 2439 |
| plant | **2389** | 2274 | 1790 |
| pig | **6622** | 4324 | na |
| *N50 RNA size (nt)* |  |  |  |
| human | 2190 | **2445** | 1358 |
| plant | 2420 | **3342** | 2395 |
| pig | 3222 | **4172** | na |
| *Express coverage-span, cover-depth* |  |  |  |
| human | 80%, 5 | 83%, 5 | **96%, 10** |
| plant | 76%, 1 | 74%, 1 | **97%, 2** |
| *TransRate coverage-span, score* |  |  |  |
| human | 34%, 0.19 | 46%, 0.24 | **87%, 0.73** |
| plant | 33%, 0.15 | 36%, 0.15 | **84%, 0.67** |

Along with filtered counts, three gene quality statistics are presented: AA1K protein sizes, N40 transcript sizes, and Expression coverage span/depth, as well as TransRate’s expression statistics are given. As expected, and shown in bold-face, maximal values of these three quality statistics coincide with the type of filter used.

There are biological maximums to protein sizes, in part because longer codes receive more random mutations and disruptions, notably the 120,000 bp long code of TITIN in human and vertebrates. Because length of proteins and protein homology are strong biological criteria, a good measure of coding gene set accuracy is one based on protein sizes. The AA1K statistic (Gilbert 2013), average of the longest 1000 proteins in a species gene set, is a biological quality measure that is simple to calculate. It correlates well with homology.

Measures of transcript sequence length are commonly published such as the N50 statistic borrowed from chromosome assembly methods. While this makes sense for chromosomes where fragments assembled are 100,000 smaller than the whole, this has little sense for transcripts that are often complete, fragments are near complete, and errors extending length are common. A frequent mistake of longest RNA methods for measuring gene quality are collection of insertion errors.

Measures of RNA-seq expression level, coverage and other qualities derived from mapping RNA reads onto transcripts are of interest. But these are technically dependent on alignment algorithms, statistical assumptions used in expression experiments, and discrepancies with transcript assembler methods. These factors can lead to mis-calculation of transcript quality, with uncertain biological interpretation for gene model quality.

There is a good correlation of conserved genes recovery with the AA1K statistic (r=0.78 to 0.91), for the transcript assemblies of vertebrates, plants and arthropods examined by Smith-Unna et al. (2016, TransRate Sup. Table 2). The TransRate score for these has a lower correlation with conserved genes (r=0.53 to 0.77). Supplemental Table TSAQ contains average transcript assembly quality scores for 155 TSA entries, adding AA1K and conserved unique protein scores to those published, and Supplemental Table AC7 has the correlation matrices of these scores.

### R: Updated EvidentialGene transcript classification

A useful result of these gene filter comparisons has been to identify problems in EvidentialGene’s implementation of its basic classifier for “longest CDS and valid alternates”. Recovery of accurate alternates and paralogs was lower than expected, and had a drop in number of valid transcripts per alt, par classes similar to “longest RNA” with CD-HIT, although overall recovery was higher. The original tr2aacds algorithm should not have caused such a drop, but instead kept valid alternates.

This author examined implementation details using these comparisons, and identified specific problems:

p1. Perfect fragment removal was removing valid alternate 5’starts, by ignoring 5’UTR sequence where alternate exons are spliced to produce shorter, but fragment-identical coding sequences. This is qualitatively different form of alternate splicing that the more common exon-alternation in CDS. The updated implementation adds 5’UTR for measure and reduction of perfect-fragment CDS alternates; UTR-only alternate splicing is still ignored by evigene methods.
p2. Transcript aberrations includes a measure of percent CDS, where transcripts with long UTR, e.g. less than 50% CDS, were reduced by that measure. However, coding genes with longer UTR are now a common occurrence in reference gene sets, increasing since 2010, as evidence of long-UTR transcript assemblies has been incorporated in reference gene sets. Now this reduction weight is only applied to transcripts with less than 20% CDS.
p3. Alternates with nearly identical proteins, as opposed to CDS, were reduced. This is a mistake based on reference gene sets, with valid CDS alternates that produce near or identical proteins.
p4. The ORF calculator option to distinguish complete versus 5’ partial proteins was adjusted for better agreement with reference gene sets, increasing portion of complete proteins, with a related result that fewer CDS alternates, at 5’ partial end, are retained.

These corrections have the general result of retaining more alternate models. New transcript set qualities have been examined and found useful additions to this “longest CDS and valid alternates” self-referential classifier:

q1. Coding potential calculation, to help distinguish random sequence that has codon chains but is less likely to be valid protein. This is common for the shorter putative CDS, where random results with large samples contain such codon chains. Various coding potential calculations are known (eg Kang et al. 2017), but are not perfect classifiers as many known reference proteins score as having no coding potential. Evigene’s implementation uses coding potential calculation as a modifier quality, to reduce low scoring transcripts in absence of other positive qualities (ie homology). A subclass ‘nc’ for non-coding potential is added to the classifications and transcript measures.
q2. Replication across assemblers of the same coding sequence transcripts is a form of technical validation, where replicated forms have higher likelihood as accurate than un-replicated alternate models (Voshall 2018). This also is not a perfect classifier. With many complex genes, a single assembly parameter/data set often produces the most accurate model. Another type of replication score, across populations or individuals of a species, can be distinguished. These are analogous to biological and technical replication approaches used in gene expression analyses.
q3. Alternate exon splicing patterns are assessed from self-alignment of gene alternates. This is an approximate measure that compares to alternate exon splicing found with chromosome mapped transcripts. Valid alternates typically share common or constitutive exons, invalid models do not share these, or contain a fragment of constitutive exons. This exon splice pattern measure helps discriminate valid and invalid models of alternates.

These updates to EvidentialGene software are available as version 4. The pipeline flow is same as described for SRA2Genes with is a complete gene reconstruction pipeline, including RNA-seq data selection, over-assemblies produced by the methods described in Gilbert (2019), the “longest CDS” reduction, followed by several external evidence measures and a 2nd stage reduction based on homology to other species, contamination checks, chromosome mapping qualities, and more.

However, a self-referential pipeline for transcriptome reduction is a valuable tool in itself, and remains a stand-alone portion of EvidentialGene. The values of this include (a) simplicity, with fewer complexities of data, no data required beyond transcripts, and fewer mistakes of omission, (b) computationally very efficient, (c) useful to compare with other gene reconstruction methods. This updated simple pipeline has two stages: at stage 1, *tr2aacds*, transcripts are classified and reduced with coding sequence metrics, then stage 2, *trclass2pubset*, this reduced set is re-classified, further reduced, and annotated for publication, with added self-referential measures noted above. This is packaged for one-step use, with input of transcriptome over-assembly, and output of a non-redundant gene-transcript sequence set, classified by loci, with quality attributes. Supplemental document EVUP has further details of these updates to gene reconstruction algorithm and implementation.

### R: Recommendations for gene set reconstruction and quality measure

Based on results with recovery of reference gene sets, and experience in this field, the author notes these problems and recommends these actions to gene information prospectors, as listed in Table 5. For measuring coding genes, “longest CDS” has the most direct biological support, and highest accuracy. “longest RNA” approaches suffer from accepting errors and measuring untranslated sequence qualities without biological support, in comparison to longest CDS. “most RNA-seq” has biological relevance to add to longest CDS measures, however implementation of this needs to distinguish technical discrepancies from expression biology.

**Table 5.** Discrepancies and recommendations for transcript classifier methods.

| Filter | Longest CDS | Longest RNA | Most RNA-seq |
| --- | --- | --- | --- |
| Overall Accuracy | highest | middle | lowest |
| *Discrepancies* |  |  |  |
| Common mistakes | longer aa | short aa, insert errors | short rna and aa, multimap failures, discrepant with denovo assembly methods |
| Unique transcripts | kept most | kept many | lost many |
| Alternates | kept most | lost many | lost many |
| Paralogs | kept most | lost many | lost many |
| *Recommendations* |  |  |  |
| Reconstruction use | primary for coding genes | not useful | expert-only, secondary or non-coding filter |
| Expertise & effort | low | low | high, TransRate errors |
| Compute resources | low | high | high |

Longest CDS measures, which result in highest accuracy and most biological meaning, are recommended for coding gene transcripts. Longest RNA measures offer no apparent added value, and result in error selection, should be avoided for coding gene transcripts. Even for non-coding transcripts, longest RNA is a quality of dubious value. Most RNA-seq measures have biological meaning, but as they can be confounded by technical effects, it is recommended they be used with care in selecting transcripts, especially when in conflict with other evidence measures.

## Discussion

### D: Value of accuracy for conserved genes

EvidentialGene will recovery accurate gene sets with few missing orthologous genes, from RNA-seq data alone, often surpassing chromosome-modeled gene sets from common sources, as indicated in Table 6 for several animals and plants. A conserved gene miss rate above 2%-5% is a correctable error in gene reconstruction. Correcting such errors leads to improved phylogenetic accuracy, as Deutekom et al. (2019) report, falsely inferred absences, averaging 18% across species examined, have effects on comparative studies. The gene reconstruction sources of Table 6 include popular and respected methods for genes modeled on chromosomes (NCBI, Ensembl, MAKER), and *de novo* transcript assemblies (Trinity short-read, PacBio long-read). These are not recovering complete conserved gene sets of well-studied species, as indicated by Evigene’s recovery rate.

**Table 6.** Conserved genes recovery in animals and plants, by EvidentialGene compared with gene sets of NCBI, Ensembl, MAKER, Trinity and PacBio methods. EvidentialGene sets are from *de novo* assemblies of RNA-seq, and comparisons are with gene sets current at the date of measure, with same available gene evidence. Numbers are percentages, At: *Arabidopsis thaliana* models recover At_Araport genes, Zm: *Zea mays* models recover sorghum, Bt: *Bemisia tabaci* models recover pea aphid, Dp: *Daphnia pulex* models recover *Daphnia magna*, Ss: *Sus scrufo* models recover human, and Dr: *Danio rerio* models recover cavefish genes. Details are in supplement table SDG9.

| Gene source | Plants |  | Arthropods |  | Vertebrates |  |
| --- | --- | --- | --- | --- | --- | --- |
|  | At_plant | Zm_corn | Bt_whitefly | Dp_wflea | Ss_pig | Dr_zfish |
| Evigene | 95% | 83% | 81% | 72% | 97% | 97% |
| NCBI |  | 81% | 80% |  | 96% | 94% |
| Ensembl/Gr |  | 82% |  |  | 95% | 93% |
| MAKER |  |  | 77% | 59% |  |  |
| Trinity | 88% |  | 73% |  |  |  |
| PacBio | 58% | 78% |  |  |  |  |

### D: Value of accuracy for alternates and paralogs

The human reference gene set of NCBI RefSeq contains 5.6 transcripts per coding locus, many more than the plant (1.7 t/l) or pig (3.0 t/l) reference sets. Yet the 113,000 transcripts of this reference set is half that found by RNA-seq assembly projects (Morillon & Gautheret 2019), with evidence of biological effects in those additional transcripts. With large effort, the human gene set is now the most accurate and complete of complex eukaryotes, implying that alternate and paralog transcripts are under-reported in most animal and plant gene sets. The results presented indicate one of the reasons: computational methodology is missing, or misclassifying many accurate transcripts. There are reasons to think that large numbers of alternates, and paralogs, are of little biological importance (e.g. Tress et al. 2017), but biological value is complex to measure. A complete set of putative alternates and paralogs facilitates the investigations into their biological value. For example, studies of venom genes (snakes: Modahl et at 2020, Holding et al. 2018, wasps), animal sense receptor genes (e.g. olfactory genes, eye pigments), plant disease resistance genes, all indicate importance of recent evolution of gene duplications in adaptation to environment.

The range of biological complexity and variation found in alternate transcripts and paralogs is high, and potential for reconstruction artifacts is similarly high, in part because both natural and artifactual causes produce the same types transcripts (eg. Dougherty et al. 2018 for cross-spliced paralogs). Long-read, single molecule sequencing methods have demonstrated value at recovering accurate transcripts, but as Morillon & Gautheret (2019) indicate, cost-effective and accurate short-read RNA data is an abundant resource for improved computational reconstruction of transcripts. The uncertainty in measurement accuracy of transcripts can be reduced with additional evidence evaluations, including replication over methods, and over biological samples.

### D: Importance of standard ORF calculations

Transcripts have the major biological quality of coding sequences and proteins. These are simple to calculate with well-established codon lookup tables, and the results match known reference proteins well. There are exceptions in reference protein agreement, as experimental evidence and expert curation finds such exceptions in standard biological translation. More complex ORF calculators are attempting two major tasks: proper translation then classification and removal of non-coding or random errors. For the examples reported in Table 2, the standard codon lookup tools are accurate, and the more complex predictors are failing at rather high rates (20%), by mis-classifying accurate ORFs. Other predictive ORF tools exist, with possibly greater accuracy (e.g. https://github.com/PacificBiosciences/ANGEL). This author’s strong recommendation is keep these tasks separate in gene reconstruction: use standard ORF calculations always for identifying most likely translation, then add coding potential and related coding measures as separate qualities. As separate qualities they can be balanced with homology to known proteins, clade-level conservation measures, conserved motifs, and others.

### D: Basic flaws reported

Basic flaws in gene reconstruction tools are identified by applying fairly simple tests against reference gene sets. That such flaws have gone unreported or unappreciated in tools that are now popular, and that are frequently used as components of many gene reconstruction pipelines, is an indication of complexity of this area, and of problems in progress of science engineering. Some of the tools examined are well-designed but mis-applied, some are well-conceived but poorly implemented. Most of the examined and related tools are more complex, and harder to check, than is useful for the particular task at hand, gene set reconstruction.

For example, EvidentialGene’s primary reduction method got out-of-sync with its intent, with respect to alternate and paralog recovery, due to detailed “improvements” (fragment detection, putative aberrations, ORF calculations) that were not as well tested against reference genes as should have been, along with a mistaken inference by this author that these improvements had only good effects. Poor side effects in complex information engineering can be measured by producers of gene information methods. The same process failure is implied for TransRate and TransDecoder, both of which can and likely will be corrected. The tests reported here are simple enough for others replicate or apply to similar uses.

### D: “longest RNA transcript” and CD-HIT qualities

CD-HIT was developed first to cluster protein sequences, then extended to short (EST) nucleotide sequences (Li and Godzik 2006). It is reliably accurate in alignment clustering, and more efficient than BLAST for similar functions. CD-HIT classifies the longest sequence in each cluster as “representative”. The flaw shown here is in applying this to gene transcripts: the longest assembled transcript in a cluster often contains errors, and discards valid primary coding sequences, as well as valid alternates and paralogs. For example, transcript ‘CCCGAGCCCACCATCGACGAG’ translates to ‘PEPTIDE’, selected by longest CDS metrics, while the longer ‘CCCtGAGCCCACCATCGACGAG’, chosen by longest RNA metrics, translates to ‘P*ahhrr’ with its insert error.

### D: “longest coding sequence” and EvidentialGene qualities

A longest CDS quality is subject to errors measuring CDS: some conserved genes have a longer CDS on reverse strand, and many have 5’UTR partial extensions. Longest CDS is a biologically useful quality, as many errors in RNA sequencing and assembly affect the protein code, and 2/3 of random errors may be avoided. While longest CDS is its primary measure, EvidentialGene methods use qualities beyond longest CDS to classify valid transcripts, as described in results.

### D: “most RNA-seq” and TransRate qualities

There is a flaw common to “most RNA-seq” measures of transcript quality, it is the interpretation that each read belongs to a single transcript. This is likely of value when measuring expression levels, but it conflicts with measuring the quality of a transcript as a biological molecule, when it shares sequence spans with other transcripts. There are variations in statistical application of this interpretation by tools focused on expression qualities. TransRate results can be dissected into components from different tools. Table 4 lists two coverage-span qualities, the first “Express coverage-span” from ‘snap’ read aligner, a 1st step in TransRate, then “TransRate coverage-span” from ‘salmon’ aligner used by TransRate at 2nd step. The second coverage score is much lower for longest-CDS and longest-RNA measures, specifically due to how salmon is assigning multi-map reads to a single transcript.

Most RNA-seq methods have technical problems. RNA-seq read mapping has uncertainties, and tools for this produce different results. More important is when these conflict with *de novo* transcript assembly using graph methods. *De novo* assemblers build transcripts with the same qualities of complete and homogeneous coverage, and read-pair accuracy and agreement, that read-map-back tools are measuring. The discrepancies between these are not resolved. Results of this report indicate that one such readmap-back method is mis-calculating expression qualities that assemblers also use, scoring low and eliminated a large portion of valid primary, alternate and paralog transcripts. Others find RNA expression measures are inconsistent with protein measures (Tress et al. 2017, Ojeda et al. 2019). Measurement of reads that map to several transcripts is not consistent among tools (eg Sahraeian et al. 2017), and degrades scores for genes with alternates and paralogs. It also appears that choice of qualities may select for shorter transcripts that contain the most common exons.

Other software and methods of measuring transcript qualities from RNA-seq may be useful in classifying accuracy of transcripts. RSEM-Det depends on having a reference transcript set, so it is unsuited for intrinsic-only evidence measures. The method of Ma and Kingsford (2019) addresses accuracy of alternate transcripts, with error estimates from observed by expected multi-map reads that may avoid problems in this area found in TransRate. Care should be taken to balance the use of expression and protein measures of gene transcript quality.

### D: Discrepancies with related work

#### “Results disagree with many reports that say best transcript assembler is Trinity”

There are many publications from this decade that compare transcript assemblers, and conclude that Trinity, or StringTie, or PacBio long-RNA, or any single assembler or modeler, has greatest accuracy, precision, or mixture of values. There are also many publications that reach other conclusions, commonly that no single assembler or reconstruction method, with single set of options, is superior for all gene loci. These reports find combinations of methods recover more accurate genes at different loci (eg Haznedaroglu et al. 2012, RNACocktail of Sahraeian et al. 2017, DRAP of Cabau et al. 2017, venom transcripomics of Holding et al. 2018). This later conclusion appears verified, as most if not all of publications supporting a single method are using a restricted set of methods, and/or technical measures that misinterpret biological gene accuracy. The conclusion that a combination of methods and options is superior arises when effort is made to test variations and measure results for each gene, against biological data sets. Average statistics of gene reconstruction methods hide a wide variation in results, where each of many thousand gene loci are independent, measurable objects. There are other considerations for gene reconstruction, such as cost/effort, expertise needed, complexity, intended uses, that may warrant certain approaches. The computational cost and expertise needed for EvidentialGene methods used here are low, indeed 2 or 3 assemblers other than Trinity, reduced to one set with Evigene, produce a more accurate gene set in less compute time, than the current Trinity assembler.

#### “Aren’t there too many genes compared to known gene sets?”

Many practitioners of gene informatics use the count of transcripts from an assembly or modeling set to assess quality. The problem with this is that one is often counting errors of omission, and mistaken models, in small transcript sets. One needs first to assess quality of each gene/transcript, with homology and other cross-species measures, with chromosomal and related same-species measures, with protein and expression self-referential measures. This will help identify accurate versus inaccurate models, where the count of accurate models increases with additional assembly sets (eg. Fig 1). When using many assemblers and options to produce over-assemblies of gene sets, one increases both the number of accurate models, and the number of ambiguous models. Ambiguous models need evidence beyond self-referential measures to re-classify. The efficiency of self-referential reduction, where it retains these many additional accurate models, allows one to apply more costly measures of data reduction with external evidence to produce gene sets with superior accuracy.

Two sub-classes of ambiguous transcripts increase with added assembly:

1. alternate transcripts that may or may not be biologically valid, the distinction needs as much added evidence as feasible. Known cases of complex genes will include 100s of biological alternates with a few, such as hyper-DSCAM in arthropods having 10,000s of measurable biological alternates. Added evidence from chromosome-mapping and splice site pattern analyses will re-classify between useful and useless alternate transcripts.
2. short, unclassified proteins, below 120aa-140aa and not alternates of longer genes, are very common and very difficult to classify between biological or random sequence results, with a majority from random sequence. These should be re-classified with homology and any species-group or experimental evidence of short proteins, eliminating a large pile of uninformative expressed sequences with some coding potential. Coding potential calculations can help, but biological, short proteins often enough are classed as non-coding by such calculations to prevent this use as an absolute classifier. Expressed transposons, where found, tend also to produce large numbers of slightly different, short proteins, and can be reclassified with transposon domain analyses. Experience of this author finds expressed transposons are generally rare, but level is species/experiment dependent.

This second sub-class is sometimes called ‘microproteins’ or small open reading frames (smORFs, Bazzini et al. 2014). There is no consensus on how to classify these transcripts in absence of experimental support, other than that further evidence, i.e., conservation across species, is desired. For human reference gene sets, NCBI RefSeq has about 2,000 while Ensembl has about 20,000 short-protein mRNA transcripts. Upstream small ORFs (uORFs) from 5’UTR of transcripts are common in vertebrates and plants (Johnstone et al. 2016, Juntawong et al. 2014), are translated and involved in gene regulation. Genes classified as non-coding or pseudogenic are also known to have expressed, translated and functional products (Ji et al. 2015). As these studies indicate, there are a wealth of expressed transcripts with small ORFs of real or potential biological importance, whether classified as coding genes or not.

In this report, the Evigene draft transcripts sets for human and plant are 4x-5x above reference set counts, almost all of the excess are unclassified short proteins. This human draft set has 341,026 putative gene loci, but only 27,240 have proteins of 140aa or longer, compared with 18,418 in the human reference set. The 313,786 short putative loci contain most of the 1,770 found in the human NCBI Ref-Seq set, and most of the spurious loci that can be discarded with further evidence classification.

The objective and repeatable measures used here for basic comparisons to known reference data, can and should be employed by scientists with interests in accurate genome information. Discrepancies between results reported by others can be examined to learn the information engineering details that cause those discrepancies. This area is complex, but an approach that compares methods and applies the simplest methodology consistent with accuracy is suitable, whether for a final product, or to compare with results from other methodology. That is the approach that EvidentialGene aims for in reconstructing accurate, reliable gene information.

An underlying theme is that popular and well-reputed tools in genomics are to be questioned with objective tests, ones that show flaws in accuracy for gene reconstruction. Partly this is because some measures in common use, such as transcript sizes and expression measures, are not the most biologically relevant ones for measuring gene accuracy. Another part of this depends on how tools are applied to gene reconstruction. Often the usage reported by authors of those tools, and default tool options, are not what works well with today’s gene data, but experimentation with options will find better ones.

Another part of these results that differs from others, is the value/cost balance used in assessing results. It is common for statistical comparisons to report false positive/negative and related scores, with an implied assumption that these are equal in weight. This is not the case for biologically relevant gene information, where accurate genes are valuable in future studies, omissions of true genes hamper those future studies. The cost to future studies of putative genes of uncertain reliability, or false positives, is low, in part because they can be ignored where evidence is lacking, in part because there is biological evidence for transcripts reconstructed from RNA expression. Positive evidence should be used to classify gene models, as to accurate or inaccurate, rather than lack of evidence. There is always more evidence to be gathered, it can be measured against unclassified transcripts, but not against omitted ones.

### D: EvidentialGene for gene reconstruction

EvidentialGene is a gene reconstruction pipeline with a demonstrated higher accuracy at recovering reference transcript sets than other popular methods for multiple transcript assemblies, and higher accuracy than any single RNA-seq assembly method. The improvements reported here for EvidentialGene, including replication and alternate exon chain analysis, make it a more robust, biologically-oriented information engineering tool. EvidentialGene is a work in progress: there are additional parts of a gene reconstruction that are partly incorporated (non-coding gene classification, contamination and error checks), and basic methods to be improved (homology annotation and classification). It now contains the needed components, fairly simple, reliable, tested and integrated, to produce gene sets that are more accurate and complete than other methods, including NCBI and Ensembl’s reference gene annotation pipelines (Gilbert 2019).

## Conclusions

The recommended approach to obtain a draft set of accurate coding genes from transcript assemblies, encapsulated in EvidentialGene methods, is (a) over-assembling with several methods and parameters, (b) measuring unique, duplicate and fragment coding sequences, (c) selecting models with longest coding sequences, modified to (d) retain alternate and paralog transcripts with sufficient unique sequence, (e) add replication of assemblies as a quality criterion, (f) add coding potential calculations to separate likely coding from non-coding or error transcripts. This results in a consistently high retention of accurate models, and reduction of extra models to a low level suited to further evidence analyses (e.g. homology to other species). In comparison, filtering large over-assemblies with longest RNA measures, as with CD-HIT-EST, and most RNA-seq expression measures, as with TransRate, discards a large portion of accurate models, especially alternates and paralogs. The EvidentialGene approach, emphasizing coding sequence quality, provides an algorithm that is consistent with gene biology and phylogenetic measures.

## Draft Notice

This is a draft paper of EvidentialGene methods. Please do offer comments and suggestions to improve this, of any sort, as it will help much in getting this across to future readers. As the paper notes, there is a substantial update to Evigene software in the wings, not yet packaged for public use. If you are willing to spend time with a still messy version for testing, let me know. The update will change transcriptome results, for the better, and if you can compare old versus new results, I especially want to hear of that. The major change is to retain more valid alternate and paralog transcripts, which prior versions are discarding too many of (but not as many as other methods). Other changes reduce spurious/redundant transcripts using added measures (described in paper). --Don Gilbert

## Acknowledgements

XSEDE/TeraGrid shared computational resources, for a decade of development and implementation, Award# MCB100147, to Genome Informatics for Animals and Plants, D.G. Gilbert.

## Supplemental Information

**Supplemental Table AC1.** Detailed statistics, including component assemblers, for accuracy at reference recovery.

**Supplemental Document OCC**. ORF computations comparison details (reference check codes, method command calls).

**Supplemental Table AAP**. Accuracy at alternate and paralog reference recovery, per filter and per assembler. Table columns are: species, filter, ngene, palt, nalt, ppar, npar, puni, nuni. Filter labels are evg9i = longest CDS with evigene, longcd = longest RNA with cd-hit-est, trate = most RNA-seq with transrate, and assemblers idba, soap, trin=trinity, velv=velvet/oases. ngene is number of accurate transcripts recovered in gene family (values of 1 to 5, uni has 1 only), palt, nalt are alternates, with p percent relative to all assemblies, and n count of accurate assemblies per filter, ppar, npar for paralogs, and puni, nuni for unique transcripts.

**Supplemental Document EVUP**: Evigene updates to self-referential gene classification and reduction algorithm (explains prior algorithm, mistakes discovered, and updated algorithm, with new code and use pipeline).

**Supplemental Table TSAQ**. Quality score table for transcript assemblies from NCBI TSA, from TransRate Sup. Table 2 (Smith-Unna et al. 2016), augmented with scores of AA1K, AAMax of Evigene, and percent complete conserved genes of BUSCO. Table columns are TSAid, tratesc, rdlen, rdpairs, tool, phyla, aa1k, aamax, n50, ntr, buscoval. TSA transcript sequences were accessed from NCBI Genbank in Nov 2016.

**Supplemental Table AC7**. TSA gene set assemblies from TransRate Sup. Table 2 (Smith-Unna et al. 2016), adding protein AA1K score and conserved genes BUSCO score. Correlations of scores, where scores are tratesc: published transrate score, rdlen, rdpairs: read length and number of paired reads, aa1k, aamax: average protein size of 1000 longest, and maximum protein size, ntr: number of transcripts in TSA assembly, busco: percent complete conserved genes, in BUSCO v9 data for vertebrates, plants and arthropods. Conserved genes score is better correlated with AA1k than TransRate score.

**Supplemental Document TRFIX**. TransRate software code patches applied by this author.

**Supplement Table SDG9.** Conserved genes recovery in animals and plants, by EvidentialGene compared with current gene sets of NCBI, Ensembl, MAKER, Trinity and PacBio methods. EvidentialGene sets are preserved for public use at persistent repository https://scholarworks.iu.edu/ as *Arabidopsis thaliana*, DOI: 10.5967/p26h-we78; *Zea mays*, DOI: 10.5967/pmst-pp68, *Bemisia tabaci*, DOI: 10.5967/wj7m-ea71, *Daphnia pulex*, DOI: 10.5967/gcs3-x192; *Sus scrufo*, DOI: 10.5967/K8DZ06G3; *Danio rerio*, DOI: 10.5967/sgv6-n273; Table from eugenes.org:/EvidentialGene/evigene/docs/evigene_plantsanimals_2017sum.txt

## Data and Software Citations

### Data sets used

*Arabidopsis* plant gene set used is Araport11_genes.201606 from xxx, accessed on 29 Jan 2017 Human reference gene set used is from NCBI at ftp.ncbi.nih.gov/genomes/refseq/vertebrate_mammalian/Homo_sapiens/reference/GCF_000001405.38_GRCh38.p12/ accessed on 24 Apr 2018.

Pig reference gene set used in this study is from NCBI at ftp://ftp.ncbi.nlm.nih.gov/refseq/S_scrofa/mRNA_Prot/pig.1.rna.gbff.gz, accessed on 27 Apr 2018.

RNA-seq sources from SRA, with NCBI BioProject ID are *human*: SRR3192658 read set of PRJ-NA30709 (ENCODE); *pig3c*: 9 read sets of PRJEB8784 (Univ. Illinois); *weed3rd1*: SRR3664432, SRR3664433, SRR3664434 read sets of PRJNA323955 (Virginia Tech.)

Vertebrate and plant conserved single-copy genes, of OrthoDB v9 (http://www.orthodb.org), BUSCO.py software, with hmmer (v3.1, http://hmmer.org/) (Simao et al. 2015).

### Software tools used

blastn, blastp of https://blast.ncbi.nlm.nih.gov/ (Altschul et al. 1990)

fastanrdb, of exonerate, https://www.ebi.ac.uk/about/vertebrate-genomics/software/exonerate (Slater & Birney 2005)

CD-HIT, cd-hit-est, of https://github.com/weizhongli/cdhit/ (Li & Godzik 2006)

EvidentialGene, of http://eugenes.org/EvidentialGene/ (Gilbert 2012) TransRate, of http://hibberdlab.com/transrate (Smith-Unna et al. 2016) velvet, oases of velvet1210 assembler, https://www.ebi.ac.uk/~zerbino/oases/ (Schulz et al. 2012) idba_tran, of idba assembler, http://hku-idba.googlecode.com/files/idba-1.1.1.tar.gz (Peng et al. 2013) SOAPdenovo-Trans, http://soap.genomics.org.cn/SOAPdenovo-Trans.html (Xie et al. 2013) Trinity, of trinityrnaseq assembler, https://github.com/trinityrnaseq/trinityrnaseq (Grabherr et al. 2011) genemark/gmst.pl and genemark_suite/gmsn.pl of http://exon.gatech.edu/GeneMark/ (Tang et al. 2015) TransDecoder.Predict and TransDecoder.LongOrfs of https://github.com/TransDecoder/ NCBI standard ORF finder of https://www.ncbi.nlm.nih.gov/orffinder/

