## Supplementary material for "Longest protein, longest transcript or most expression, for accurate gene reconstruction of transcriptomes?": supAC7_tsa_trqual_correlations.pdf

**Table AC7**. TSA gene set assemblies from Transrate paper table XXX, adding protein AA1k score and conserved genes BUSCO score. Correlations of scores, in Table SAC7, where scores are tratesc: published transrate score, rdlen, rdpairs: read length and number of paired reads, aa1k, aamax: average protein size of 1000 longest, and maximum protein size, ntr: number of transcripts in TSA assembly, busco: percent complete conserved genes, in BUSCO v9 data for vertebrates, plants and arthropods. Conserved genes score is better correlated with AA1k than Transrate score.

*Vertebrates (n=35)*

| *Corr* | tratesc | rdlen | rdpairs | aa1k | aamax | ntr | busco |
|---|---|---|---|---|---|---|---|
| tratesc | 1 | 0.21 | -0.19 | 0.43 | 0.53 | -0.50 | 0.70 |
| rdlen | 0.21 | 1 | 0.53 | 0.58 | 0.32 | 0.34 | 0.58 |
| rdpairs | -0.19 | 0.53 | 1 | 0.41 | 0.13 | 0.52 | 0.24 |
| aa1k | 0.43 | 0.58 | 0.41 | 1 | 0.55 | -0.04 | **0.91** |
| aamax | 0.53 | 0.32 | 0.13 | 0.55 | 1 | -0.26 | 0.63 |
| ntr | -0.50 | 0.34 | 0.52 | -0.04 | -0.26 | 1 | -0.26 |
| busco | 0.70 | 0.58 | 0.24 | 0.91 | 0.63 | -0.26 | 1 |
| *Means* | 24 | 89 | 35M | 1344 | 5080 | 79181 | 52 |

*Plants (n=28)*

| *Corr* | tratesc | rdlen | rdpairs | aa1k | aamax | ntr | busco |
|---|---|---|---|---|---|---|---|
| tratesc | 1 | -0.29 | 0.04 | 0.66 | 0.56 | -0.07 | 0.77 |
| rdlen | -0.29 | 1 | 0.21 | 0.16 | 0.13 | 0.01 | 0.06 |
| rdpairs | 0.04 | 0.21 | 1 | 0.05 | 0.03 | 0.19 | 0.11 |
| aa1k | 0.66 | 0.16 | 0.05 | 1 | 0.76 | -0.01 | **0.78** |
| aamax | 0.56 | 0.13 | 0.03 | 0.76 | 1 | -0.18 | 0.65 |
| ntr | -0.07 | 0.01 | 0.19 | -0.01 | -0.18 | 1 | 0.12 |
| busco | 0.77 | 0.06 | 0.11 | 0.78 | 0.65 | 0.12 | 1 |
| *Means* | 24 | 90 | 43M | 932 | 2963 | 57551 | 49 |

*Arthropods (n=58)*

| *Corr* | tratesc | rdlen | rdpairs | aa1k | aamax | ntr | busco |
|---|---|---|---|---|---|---|---|
| tratesc | 1 | 0.09 | -0.31 | 0.34 | 0.42 | -0.03 | 0.53 |
| rdlen | 0.09 | 1 | 0.20 | 0.28 | 0.31 | 0.29 | 0.17 |
| rdpairs | -0.31 | 0.20 | 1 | -0.25 | -0.16 | 0.09 | -0.17 |
| aa1k | 0.34 | 0.28 | -0.25 | 1 | 0.90 | 0.38 | **0.78** |
| aamax | 0.42 | 0.31 | -0.16 | 0.90 | 1 | 0.29 | 0.78 |
| ntr | -0.03 | 0.29 | 0.09 | 0.38 | 0.29 | 1 | 0.31 |
| busco | 0.53 | 0.17 | -0.17 | 0.78 | 0.78 | 0.31 | 1 |
| *Means* | 18 | 95 | 34M | 1098 | 4135 | 41353 | 63 |
