## Supplementary material for "Longest protein, longest transcript or most expression, for accurate gene reconstruction of transcriptomes?": supDG9_evigene_plantsanimals_2017.txt

Animal and Plant gene set reconstructions with EvidentialGene
Comparisons to other popular and recent gene reconstructions
In-progress, 2016/2017, D.G. Gilbert
-------------------------------------------------

Tables summarizing alignment of test gene sets to reference species
  1. Arabidopsis model plant
  2. Corn (Zea mays) crop plant
  3. White fly (Bemisia tabaci) crop plant pest 
  4. Water flea (Daphnia pulex) environmental sentinal species
  5. Zebrafish  gene reconstructions, Danio rerio (model organism)
  6. Pig, Sus scrufa, gene reconstructions compared
-------------------------------------------------

1. Plant model Arabidopsis thal. gene reconstruction methods compared
    Gene assemblies of Illumina RNA-seq vs PacBio, 2017.feb, d.g.gilbert

1a.  Gene assembly methods measured against unique coding sequences of 
     reference Arabidopsis public gene set (2016, At16Ap, transcripts n=37806) 

  Geneset  nFound  Found%  AlignF% AlignT% 
  At5EVm   36072   95.4    95.7    95.0     
  At2EVm   34294   90.7    94.0    92.3   
  Oases    34030   90.0    93.5    91.2   
  IDBAtr   33837   89.5    92.0    89.1   
  SOAPtr   33598   88.9    90.5    87.0   
  Trinity  33417   88.4    87.9    84.1   
  PacBio   21964   58.1    76.7    48.2   
           ---------------------------

1b. Arabidopsis gene sets measured against related species Orange and Cacao.

               Cacao Reference                Orange Reference
 Geneset  nGene  nAlt    Found% AlignF%   nGene  nAlt   Found% AlignF% 
 At16Ap   22334  131865  88.7   70.7      16795  22280  91.4   74.7   
 At17EVm5 22581  132507  89.1   70.3      16613  22087  90.6   74.2   
 At16Pacb 17593  95437   64.2   60.5      11578  15608  64.0   63.5   
          ----------------------------    ---------------------------

1c. Intron recovery for Arabidopsis gene sets (ni=125481 of RNA-seq mapped to chrs)
  Geneset  GeneTr  valExon Found%
  At16Ap    42211  110654  88.1  
  At17EVm5 141670  109859  87.5   
  At16Pacb  48848   70719  56.3   
  .. subset gene assemblies ..
  Oases    312817  101784  81.1
  IDBAtr   248429  101357  80.7
  SOAPtr    96533   99274  79.1
  Trinity  198380  102208  81.4
          ----------------------

Arabidopsis gene set versions
  At16Ap  = public gene set of 2016 of Arabidopsis thal. from Araport.org 
  Oases   = velvet/oases assembly of Illumina RNA,
  IDBAtr  = idba_tran asm of Ill. RNA,
  SOAPtr  = SOAP-Trans asm of Ill. RNA,
  Trinity = Trinity asm of Ill. RNA,
  At16Pacb/PacBio = Pac-Bio "no-assembly" assembly (PacBio xxx method) of Pac-Bio RNA data
  At2EVm = Evigene classified gene set of Illumina RNA assemblies above (O,I,S,T)
  At17EVm5/At5EVm = Evigene reduction of assemblies of Illumina RNA-seq, 
           improved over At2EVm with added RNA sets and gene assemblies
  At17EVm5 genes of 2017 are [soon to be] public at 
    http://arthropods.eugenes.org/EvidentialGene/plants/arabidopsis/evigene2017_arabidopsis/

Method: BLASTn -query reference-unique.cds -db allgenesets.cds -evalue 1e-5 ..
Statistics
  Found% = percent of reference transcripts found
  AlignF% = align to reference transcripts found.  
  AlignT% = align to total reference transcripts.  
Intron Statistics and Methods, see below 

Reference genes:
  Orange, NCBI genomes/refseq/plant/Citrus_clementina/GCF_000493195.1_Citrus_clementina_v1.0
  Cacao, Evigene update of Theobroma cacao from RNA-seq supplied by Mars.

=============================================================

2.  Corn genes reconstructions 

2a. Zea mays gene sets measured against reference proteins of Sorghum and Arabidopsis
                Sorghum proteins            Arabidopsis proteins
Geneset nFound Found% AlignF% AlignT%   nFound Found% AlignF% AlignT% 
ZmEvig  28352   82.9   92.9    91.1     22727   79.5   86.7    85.6   
ZmGram  28019   81.9   93.2    90.3     22661   79.2   86.2    84.8   
ZmNCBI  27809   81.3   93.1    89.6     22640   79.2   86.2    84.7   
ZmPacb  26680   78.0   89.3    82.4     22165   77.5   85.1    81.9   
ZmJgi4  26557   77.6   88.4    81.2     21972   76.8   85.2    81.3   
        ---------------------------     ---------------------------

2b. Restricted comparison to gene loci found in common over gene sets.
           Sorghum reference exons     Arabidopsis reference exons
Geneset   nGene nAlt   Found%  AlignF%  nGene nAlt  Found% AlignF% 
ZmEvig    16905 17255  94.5    98.2     7472  7630  81.6   74.8   
ZmGram    16905 17126  93.7    98.1     7472  7612  80.7   74.8   
ZmPacb    16905 17194  90.2    97.2     7472  7617  80.4   74.7   
ZmNCBI    16905 17046  91.6    98.0     7472  7538  78.5   74.6   
          -------------------------    ---------------------------

2c. Intron recovery for Zea mays gene sets (ni=198622 of RNA-seq mapped to chrs)
 Geneset  GeneTr  valExon Found%
 ZmEvig   171197  136609   68.7 
 ZmGram   143193  135333   68.1 
 ZmPacb   392946  135466   68.2   
 ZmJgi4    75241  136901   68.9
 ZmNCBI    na       na      na  
          ----------------------

Corn gene sets
  ZmEvig = Evigene Zeamay5fEVm 2016 assembly of Illumina RNA-seq, public at
     http://arthropods.eugenes.org/EvidentialGene/plants/corn/evg5corn/
  ZmGram = Ensembl/Gramene 2016.09 Zm000nnnn, 
  ZmPacb = CSHL/Gramene PacBio gene assemblies of 2016 as SRA entries SRR3147024..054,
  ZmNCBI = NCBI 2014 refgen zeamay
  ZmJgi4 = JGI Rnnotator assembly set of Illumina RNA-Seq , 2014
Reference genes:
  Sorghum = Sorghum bicolor 2016, Sbicolor_313, from JGI/Phytozome.org, total primary isoforms  = 34211
  Arabidopsis  = Arabidopsis thal. public gene set of 2015 from Araport.org, total primary isoforms = 28598
Method, 2a: BLASTp -query ref_proteins -db allgenesets_proteins -evalue 1e-5 ..
Statistics, 2a (more at end)
  Found% = percent of reference transcripts found
  AlignF% = align to reference transcripts found.  
  AlignT% = align to total reference transcripts.  
Method, 2b: BLASTn -query reference-unique.cds -db allgenesets.cds -evalue 1e-5 ..
Statistics, 2b
  nAlt  = number of reference alternate transcripts found, for nGene reference loci,
        restricted to reference loci common to all gene sets.
  Found% = percent of reference exons found, of total ref unique exons
  AlignF% = align to ref exons found, as percent of ref exon sizes.  
Method and Statistics, 2c : see below

=============================================================
  
3. White fly Bemisia tabaci (cotton/crop plant pest)

      Gene sets of White fly measured against reference proteins         
  Reference   Pea aphid          Fruit fly        Conserved F.Fly
Geneset  Found% AlnF% AlnT%  Found% AlnF% AlnT%  Found% AlnF% AlnT% 
BtEvig    81.2  90.0  88.0    74.1  81.8  74.9    98.7  85.4  80.6
BtNCBI    79.7  85.5  82.3    73.4  78.7  71.6    98.5  83.0  78.1
BtMakr    77.4  80.2  73.8    72.1  74.9  66.0    97.1  79.2  72.9
BtTrin    73.5  71.0  59.2    68.0  67.4  53.2    95.5  74.4  64.1
          ----------------    -----------------   -----------------

3b. Intron recovery for Bemisia tabaci gene sets (ni=134153 of RNA-seq mapped to chrs)
 Geneset  GeneTr  valExon Found% 
 BtEvig   63368   91946   68.5   
 BtNCBI   20506   93157   69.4   
 BtMakr   13825   77455   57.7   
 BtTrin   20534   67879   50.5     
          -----------------------
          
Bemisia tabaci gene sets compared
  BtEvig = Evigene gene assembly, 2016 update (vers 3), available [soon] at
    http://arthropods.eugenes.org/EvidentialGene/arthropods/whitefly/whitefly3evigene/
  BtNCBI = NCBI RefSeq gene models, 2016
  BtMakr = Whitefly genome project genes modeled with MAKER, 2016, whiteflygenomics.org
  BtTrin = TSA.GBII gene assembly 2015, Trinity of Illumina
Reference genes:
  Pea aphid Acyr. pisum, NCBI RefSeq 2016, total primary isoforms n=18601
  Fruit fly, NCBI RefSeq 2015, total primary isoforms n=13828
  Conserved Fruit fly, NCBI RefSeq 1-copy genes identified by BUSCO, total n=3055
Method: BLASTp -query ref_proteins -db allgenesets_proteins -evalue 1e-5 ..
Statistics
  Found% = percent of reference proteins w/ signif. alignment
  AlnF,AlignF% = average percent align to found reference protein 
  AlnT,AlignT% = percent align to all reference proteins 
          
=============================================================
  
4. Water flea Daphnia pulex gene sets compared

Reference  Daphnia magna        Fruit fly        Conserved F.Fly
           DpEvig7  DpMaker7  DpEvig7  DpMaker7  DpEvig7  DpMaker7
  Found%     72.0    58.9       67.9    64.3       94.9    90.3
  AlignF%    88.8    85.6       80.7    79.1       83.7    81.9
  AlignT%    88.6    69.9       80.3    74.5       83.5    77.7
  Tiny%       0.2    23.8        0.8     6.5          na
  Best%      75.4     3.1       65.4     5.2          na
           equal  21.3        equal  29.2        
  ------------------------    ---------------     ----------------

4b. Intron recovery for Daphnia pulex gene sets (ni=189710 of RNA-seq mapped to chrs)
 Geneset  GeneTr  valExon Found%
 DpEvig7   115051  126417  66.6 
 DpMaker7   18296   88747  46.7 
          ----------------------

Daphnia pulex gene sets      
  DpEvig7 Evigene genes of 2017 from 
    http://arthropods.eugenes.org/EvidentialGene/daphnia/daphnia_pulex/daphnia_pulex_genes2017/
  DpMaker7 genes of 2017 from report of doi:10.1534/g3.116.038638 
Reference genes:
  Daphnia magna water flea,  Evigene 2015 set, primary isoforms n=29127
  Drosophila melanogaster fruit fly, NCBI RefSeq 2015, total primary isoforms n=13828
  Conserved Fruit fly, NCBI RefSeq 1-copy genes identified by BUSCO, total n=3055
Methods: BLASTP -query reference.aa -db dapplx_twogenesets.aa -evalue 1e-5
Statistics: 
  Found  = percent of reference genes with signif. align in target gene set
  AlignF = average % alignment to found reference genes (align-aa/ref-aa)
  Tiny   = target genes with size < 50% of reference length, of found genes
  Best   = which target set has longest alignment per ref gene, of found genes

=============================================================

5. Zebrafish  gene reconstructions, Danio rerio (model organism)

    Reference fish gene proteins
    Reference:   Cavefish            Carp         
  Geneset   Found% AlnT% Frag% Found% AlnT% Frag% 
 -------------------------------------------------  
  DrEvigene   97.0  96.5  0.5   92.8  94.1  0.6   
  DrNCBI      93.9  92.7  3.9   85.6  86.3  8.9    
  DrEnsEMBL   93.1  90.3  5.7   85.9  84.6  9.9   
 -------------------------------------------------

  5h. Human genes represented in Zebrafish  gene sets
  Geneset    Found% AlnT% Frag% 
  ------------------------------
  DrEvigene   87.5   90.8   0.5  
  DrNCBI      86.9   89.3   1.2  
  DrEnsEMBL   86.3   88.4   2.2  
  -------------------------------

5b. Zebrafish  gene reconstructions, 
  Conserved Vertebrate genes, BUSCO n=2500
  Geneset     Align  Miss Frag
 ------------------------------
  DrEvigene   446.8    9    5
  DrNCBI      434.6   19   13
  DrEnsEMBL   428.2   29   47 
  DrTSAGFIL   435.0   57   23
 ------------------------------

5c. Zebrafish RNA introns  
  Geneset     Found% 
 --------------------
  DrEvigene    81.6  
  DrNCBI       76.4  
  DrEnsEMBL    57.6  
 --------------------


Danio rerio gene sets compared
  DrEvigene = Evigene gene assembly, 2017 Dec
    http://eugenes.org/EvidentialGene/vertebrates/zebrafish/zebrafish17evigene/
  DrNCBI = NCBI RefSeq gene models, 2016 Dec, accession GCF_000002035.5_GRCz10
  DrEnsEMBL = Ensembl gene  models, 2017 Nov
  DrTSA = TSA.GFIL, 2017 bioproject PRJNA255848 from Phylofish, INRA, PMID:27189481
  
Reference species genes for Danio rerio 
  Cavefish = Sinocyclocheilus_anshuiensis, n=28811 genes, NCBI RefSeq of 2016
  Carp = Cyprinus_carpio, n=36674 genes of NCBI RefSeq 2016
  Catfish = Ictalurus_punctatus, NCBI 2016
  Human = Homo_sapiens, n=20191 loci of NCBI RefSeq 2018
  Conserved 1 copy vertebrate genes n=2586, of OrthoDB v9, BUSCO
  RNA-introns n=316113 consensus introns from RNA-seq mapped to GRCz10 chromosome assembly

=============================================================


6. Pig, Sus scrufa, gene reconstructions compared

                Human genes      Vertebrate_BUSCO 
  Geneset    Align% Miss% Frag%  Align  Miss Frag  
  SsEvigene   97.0   0.7  1.4    447    8    10  
  SsNCBI      96.0   0.7  0.7    440   17     2  
  SsEnsembl   95.0   0.9  1.1    431   14    20  
 ----------------------------- ----------------- 

  6b. Pig gene reconstruction method subsets compared
  Method	Miss	Frag	Short	
  Velvet	 5%	 7%	23%	
  Idba   	 8%	12%	30%	
  Pacbio        12%	15%	33%
  Soap   	12%	16%	36%	
  Trinity	20%	28%	49%	
  ---------------------------------

Pig gene sets
  SsEvigene = http://eugenes.org/EvidentialGene/vertebrates/pig/pig18evigene/
  SsNCBI = NCBI RefSeq genes for the pig, 2018-May
  SsEnsembl = Ensembl genes for the pig, 2018-June
  Human RefSeq 2018 genes, 37,868 proteins matched, of 19122/20011 loci matched
  Vertebrate BUSCO set of 2586 proteins of OrthoDB v9

=============================================================

7. Green Sea Urchin L.v. gene reconstructions compared

            P. Sea Urchin   Metazoa_BUSCO 
  Geneset   Found%  AlnT%  Align Miss Frag  
  LvEvigene  xx.x   xx.x    xxx     3    1
  LvMaker    xx.x   xx.x    xxx   345  186
  PsNCBI     --     --      xxx    12   18
 ----------------------------------------------

Green Sea Urchin gene sets
  LvEvigene = http://eugenes.org/EvidentialGene/inverts/sea_urchin/
  LvMaker = http://www.echinobase.org/Echinobase/Lv_LVA_Genes
  PsNCBI  = NCBI gene set for purple sea urchin 
  Metazoa BUSCO set of 978 proteins of OrthoDB v9

=============================================================

Common methods and statistics
  BLASTp|BLASTn -query reference.sequences -db allgenesets.sequences -evalue 1e-5 

  Tabulate largest alignement of each target set to each reference sequence (targets matching 2+ refs allowed), 
  count and sum significant alignments per gene set, including 
    Found%  = found alignments, 100 * n-aligned/n-ref; 
    AlignF% = sum(100*align-bases/ref-bases)/nFound, average of per-ref percent align, for found alignments;
    AlignT% = 100*sum(aligned-bases)/sum(ref-bases)
    Tiny%   = 100 * count(n-target-size < 50% of ref-size) / nFound, for aligned target 
    Best%   = pairwise measure for 2 gene sets only, counts of target-A <=> target-B alignment per reference
          bestA = sum(alignA > minDiff + alignB)/nFound(A or B), minDiff = 9 or other
          bestB = sum(alignB > minDiff + alignA)/nFound(A or B)
          equal = sum(abs(alignA - alignB) <= minDiff)/nFound(A or B)

Two independent quality measures of gene sets are (a) reference genes found or reconstructed in some fashion, 
(b) completeness of reconstruction of reference genes (full alignment with target length =~ reference length).
AlignF is this (b) independent measure from (a) Found genes, while AlignT is roughly the union of these. 
There is a subtle bias in AlignF: those target gene sets with fewer found genes tend to
find the longer, better aligned reference genes.  AlignF can be larger where fewer are found,
even when alignments are identical between target sets for found genes.  Pairwise comparisons of 2 gene
sets, i.e. the Best metric, is not biased by the different reference subsets found between gene sets.

Additional measures of target gene sizes vs reference size, independent of alignment size, are useful;
In practice, small outliers Tiny%, indicate fragmented gene prevelance and can help distinguish gene 
set qualities. Large outliers, indicative of some kinds of joined-gene artifacts, don't usually 
discriminate gene sets, but may depending on how a gene set was constructed.

Intron Methods and Statistics  
  map RNA-seq (Illumina) to chromosome assembly with GSNAP, 
  extract splice-mapped reads and their intron locations, 
  tabulate gene-exon x rna-intron matches.
Statistics
  GeneTr  = gene transcripts total in gene set
  valExon = gene exons w/ validated intron
  InFound% = percent of all valid introns recovered b/n gene exons
          
=============================================================

    
