## Supplementary material for "Longest protein, longest transcript or most expression, for accurate gene reconstruction of transcriptomes?": supEVUP_evigene_upd1908.txt

# supEVUP_evigene_upd1908.txt

=item Updated EvidentialGene transcript classification

A useful result of these gene filter comparisons has been to identify
problems in EvidentialGene's implementation of its basic classifier
for "longest CDS and valid alternates".   Recovery of accurate
alternates and paralogs was lower than expected, and had a drop in
number of valid transcripts per alt, par classes similar to "longest
RNA" with CD-HIT, although overall recovery was higher.  The original
tr2aacds algorithm should not have caused such a drop, but instead
kept valid alternates.

This author examined implementation details using these comparisons,
and identified specific problems:
  p1. Perfect fragment removal was removing valid alternate 5'starts,
  by ignoring 5'UTR sequence where alternate exons are spliced to
  produce shorter, but fragment-identical coding sequences.  This is
  qualitatively different form of alternate splicing that the more
  common exon-alternation in CDS.  The updated implementation adds
  5'UTR for measure and reduction of perfect-fragment CDS alternates;
  UTR-only alternate splicing is still ignored by evigene methods.
  
  p2. Transcript aberrations includes a measure of percent CDS, where
  transcripts with long UTR, e.g. less than 50% CDS, were reduced by
  that measure.  However, coding genes with longer UTR are now a
  common occurrence in reference gene sets, increasing since 2010, as
  evidence of long-UTR transcript assemblies has been incorporated in
  reference gene sets.  Now this reduction weight is only applied to
  transcripts with less than 20% CDS.
  
  p3. Alternates with nearly identical proteins, as opposed to CDS,
  were reduced.  This is a mistake based on reference gene sets, with
  valid CDS alternates that produce near or identical proteins.
  
  p4. The ORF calculator option to distinguish complete versus 5'
  partial proteins was adjusted for better agreement with reference
  gene sets, increasing portion of complete proteins, with a related
  result that fewer CDS alternates, at 5' partial end, are retained.

These corrections have the general result of retaining more alternate
models.  New transcript set qualities have been examined and found
useful additions to this "longest CDS and valid alternates"
self-referential classifier:
  q1. Coding potential calculation, to help distinguish random
  sequence that has codon chains but is less likely to be valid
  protein.  This is common for the shorter putative CDS, where random
  results with large samples contain such codon chains.  Various
  coding potential calculations are known (eg Kang et al 2017), but
  are not perfect classifiers as many known reference proteins score
  as having no coding potential.  Evigene's implementation uses coding
  potential calculation as a modifier quality, to reduce low scoring
  transcripts in absence of other positive qualities (ie homology).  A
  subclass 'nc' for non-coding potential is added to the
  classifications and transcript measures.
  
  q2. Replication by assemblers of the same coding sequence
  transcripts is a form of technical validation, where replicated
  forms have higher likelihood as accurate than un-replicated
  alternate models [cite Voshall thesis].  This also is not a perfect
  classifier, as with many complex genes, a single assembly
  parameter/data set often produces the most accurate model.  Another
  type of replication score, across populations or individuals of a
  species, will be distinguished, analogous to biological and
  technical replication approaches used in gene expression analyses.
  
  q3. Alternate exon splicing patterns are assessed from
  self-alignment of gene alternates. This is an approximate measure
  that compares to alternate exon splicing found with chromosome
  mapped transcripts.  Valid alternates typically share common or
  constitutive exons, invalid models do not share these, or contain a
  fragment of the shared exons.  This exon splice pattern measure
  helps discriminate valid and invalid models of alternates.

These updates to EvidentialGene software are [soon to be] available,
as version 4.  The pipeline flow is same as described for SRA2Genes
with is a complete gene reconstruction pipeline, including RNA-seq
data selection, over-assemblies produced by the methods described in
Gilbert (2019), the "longest CDS" reduction, followed by several
external evidence measures and a 2nd stage reduction based on homology
to other species, contamination checks, chromosome mapping qualities,
and more.

However, a self-referential pipeline for transcript reduction is a
valuable tool in itself, and remains a stand-alone portion of
EvidentialGene.  The values of this include (a) simplicity, with fewer
complexities of data, and no data required beyond transcripts,  and
fewer mistakes of omission, (b) computationally very efficient, (c)
useful to compare with other gene reconstruction methods.  The updated
"simple" pipeline now requires two stages, stage 1, tr2aacds:
transcripts to classified, reduced proteins+coding sequences, then
stage 2, trclass2pubset: re-classified and reduced for publication,
with the added self-referential measurements noted above.  This can be
packaged for one-step use, input: transcript over-assembly, output:
reduced gene-transcript sequence set, with quality attributes. 
Supplemental document EVUP has further details of these updates to
gene reconstruction algorithm and implementation.

#====================================================================================
# evigene_upd1908.info

=item UPD1908 cdna_bestorf.pl
  
  * added -codepot option, coding potential classifier, hexamer loglikely code/nocode table, 
    sort of works but misclasses true cds, as others do.
  * other bestorf updates .. -report improved orf stats
  
=item UPD1908 of evigene tr2aacds2c.pl and asmrna_dupfilter3c.pl

  * improved best perfdup and perffrag choices, tr2aacds2c nrcheckaaqual,
    ie no longer picks shortest utr of cds-dups, but median utr size
  
  * replicate/agree/consensus evidence update, doesnt rescue frag/partial for agree>1  

  * bestorf ORF_FULLvPART default 0.85 should be reduced to 0.50 or lower for accurate uses,
    ref gene sets have few aapartials due largely to rarely calling long partial vs short full orfs.
  
  * aaqual table now includes Code/Noncode column from bestorf codepot qual.
  
  
  * added 'nc' class tag to tr2aacds classes, for pot. noncoding, 
    ignores 'noncode' tag if aasize >= 140, or aasize > 99 and not poor 
    keeps all 'nc' subclass of main/alt/noclass, but drops nc part/frag/utrorf,
    
  * modified default parameters: AADUP  99% from 98%, 

=item UPD1909 run_s07p_tr2cdspub.sh

  see below update tests
  run_s07p_tr2cdspub.sh encapsulates new evigene self-ref algorithm
  using updates in code and parameters:
    prot/tr2aacds2d.pl  : rename? to tr2aacds4.pl, w/ options  -aconsensus -runsteps=noaadup
    cds_bestorfxx.pl w/ options ORF_FULLvPART=0.50, CDSXUTR=900,60
    genes/blasttrset2exons2.pl : new algorithm
    rnaseq/asmrna_altreclass3c.pl : new indata, new options, reclass and rewrite okayset to publicset; 
    genes/trclass2pubset.pl    : integrate changes from run_s07p_tr2cdspub.sh ??
  
=item UPD1908 classifytr drop valid alts problem
  
    drops of 'true' reference genes, human, plant, are mainly 'althi' with 2 pflag qualities:
      aadup (>= 99% ident from cdhit.aa cluster)
      utrpoor/bad
    large portion have pia = 100/100/ but signif num have 100/99..80..small %align-global, should treat those as valids?  
    maybe drop aadup criteria, or at least balance it w/ other rescue criteria: pi/pa < 100/100 ? 
    add criteria of total non-ident cds-bases? some threshold (10? 100?) keeps alt?

    * ignoring aadup & utrpoor qualities would boost okayset by large amount (majority of drops),
      need further criteria to adjust these ..
  
    pflag, see above constants 2=utrbad 4=utrpoor 8=aadup
    see also FIXME 201402 drop.mains problem
      AAPART cutoff is problem, with UTRBAD/POOR, causing drop.mains of uniq orthologs 
      AAPART & AAMIN, AAMINBAD, AAMINPOO  need fix to prevent drop uniq/valid main/noclass genes, 
  
    drop.parthi has similar loss of valid ref alts, adjust parthi criteria?
      frag/part == $tsize < $ALTFRAG*$qsize; ALTFRAG = 0.50 default, should retest
      
    drop valid alts via perffrag: this happens when true alt is perfect substring of longer alt,
      not sure this is likely alternating-exon gene structure, but may be,
      more likely is experimentally found alternate start site in same exon structure.
      
    for ref_arath16ap.trclass
      5167 drop.althi, 3695 have aadup, 1445/1472 have utrpoor/bad of no-aadup
      5601 okay.althi, 0 have aadup, 1062 have utrpoor/bad

    for bevgc_evg3weed1rd.trclass
      85208 drop.althi, 41846 have aadup, 30681/43362 have utrpoor/bad, 14032/43362 aapartial of no-aadup
      106526 okay.althi, 0 have aadup, 32274 have utrpoorbad, 22851 aapartial

    for ref_human.trclass
      26715 drop.althi, 14346 aadup, 12305/12369 have utrpoor/bad of no-aadup
      22563 okay.althi, 0 aadup, 5975 have utrpoor/bad
    
    for trev19human.trclass
      126070 drop.althi, 45301 aadup, 37126/80769 utrpoorbad of noaadup, 35811/80769 aapartial of noaadup
      138448 okay.althi, 0 aadup, 39146 utrpoorbad, 30744 aapartial

=item update tests

  details:  evg_bestgenefilt/tr2aacds_test2ae.info
  evigenes=$HOME/bio/evigene19sep03/scripts
  evapp=$evigenes/prot/tr2aacds2d.pl
  export ORF_FULLvPART=0.50
  export CDSXUTR=900,60
  export pCDSOK=20 pCDSBAD=20 
  addopt=" -aconsensus -runsteps=noaadup"
  with tr2aacds2c.pl

#---------------------------------------------
      

=item evigene-4upd update details for pig3c example

------------------------
evigene-4upd == tr2aacds_test1908f/try9pig3c/pig3c.trclass.sum.txt
-- compare to published evg17pig3c/pig3c.trclass, both okayset and publicset
-- use to explain upd changes to evigene, tr2aacds + tr2genes (?), version 4?

Changes to evg tr2aacds v4:
-- more complete proteins, cdna_bestorf option ORF_FULLvPART=0.50 from
   0.85, to better match standard transcript/protein results;
   2ndary result is to drop more perfect duplicate coding sequences.
-- added 'nc' subclass of poor/non coding potential transcripts, method added to cdna_bestorf
-- removed 'a2' subclass & test of protein duplicate drops, as it removes valid alternates
-- tr classifier: reduced cost of long UTR (ie. aberrant or joined-gene score) as such are now a common
   occurence in reference gene sets (due to more evidence of long-UTR in transcript assemblies); 
   params pCDSOK=pCDSBAD=20 (min % of mRNA as CDS is okay quality)
-- added assembly replication quality score, keeps alternates and noclass when assemblers agree.
-- fewer perfect-fragments; modified perfect-fragment test to include 5'UTR in test of identical CDS fragments, 
  as this test was dropping most valid alternate start exons with splices in 5'UTR (proper name is what?)
-- modified keep/drop settings, esp. keep more althi, high identity alternates as these are measured 
  to contain many biologically valid alternates.
  
-- overall, many more transcripts are retained in okayset as potentially valid, but wanting further
   evidence tests to reduce to less redundant set.  This 2nd stage reduction filter is done now in
   trclass2pubset.pl, with two variants: self-referential only (using only input trasm sequences),
   and adding external evidence of more complete SRA2Genes/Tr2Genes, with cross-species homology,
   contaminant checks, chromosome map checks.
-- in keeping value of  self-referential only reduction filter, this 2nd stage reduction should always
  be used w/ tr2aacds, 
-- 2nd stage reduction filter adds new self-ref quality algorithm (genes/blasttrset2exons): 
  alternate/paralog exon splice site measurement, w/o chromosomes,
  by alt transcript alignments per gene locus, assessment of common/constituative exons shared across
  alternates, and quality scoring of alternates as to common/unique exon splice patterns.  This is
  an analog of chromosome-mapped exon splice pattern analysis.  It agrees with chr-map results enough to be 
  useful to help decide if alternate models are redundant/fragment models, 
  as tested with reference and assembled genes.
-- recommended result data are now the publicset/ portion of EvidentialGene's pipeline,
  in common with fuller SRA2Genes pipeline, with or without added external gene evidence.
  Primery results table, trname.pubids, re-classifies all transcripts of the okayset from tr2aacds,
  as to keep or cull.  Cull is a new transcript category of insufficient evidence, which can be
  either ignored or tested further for useful transcript models.


## check weed as example for class counts?
## check/recheck reclass by tr x tr self align, cds self misses utr aligns.. esp for tiny aa set
    
New, tr2aacds version 4                                 Prior, tr2aacds version 3 (published in pig pp)
==> try9pig3c/pig3c.trclass.sum.txt <==                 ==> ../evg17pig3c/pig3c.trclass.sum.txt <==
# Class Table for pig3c.trclass                         # Class Table for pig3c.trclass 
class           %okay   %drop   okay    drop            class           %okay   %drop   okay    drop
althi           1.5     0.5     135691  46302           althi           1.7     2.3     140456  189820
althi1          4.5     0.9     395244  85085           althi1          1       4       89973   334160
althinc         0.8     0       77068   0               althia2         0       0.12    0       10696
altmfrag        0.07    0.09    6647    8537            altmfrag        0.05    0.14    4474    11586
altmid          0.07    0.06    6221    5950            altmfraga2      0       0       119     256
main            1.5     0.26    134411  22820           altmid          0.07    0.13    5975    11483
mainnc          1       0       94484   0               altmida2        0       0       185     262
noclass         2.7     1.8     239579  156876          main            1.6     1.4     138765  116931
noclassnc       1.7     0       154640  0               maina2          0.08    0.02    7130    1761
parthi          0       1.6     0       144831          noclass         2.1     3.8     177421  315510
parthi1         0       1.2     0       104864          noclassa2       0       0       25      67
perfdupl        0       65.1    0       5655186         parthi          0       2       0       169901
perffrag        0       13.8    0       1205789         parthi1         0       0.35    0       29104
---------------------------------------------           parthia2        0       0.04    0       4119
total           14.3    85.6    1243985 7436240         perfdupl        0       59.8    0       4941776
=============================================           perffrag        0       18.7    0       1549765
okay.top n=1000; ave=2130; min,max=1385,31655; full=983;     ---------------------------------------------
okay.all n=623114; ave=73; min,max=16,31655; full=547822;     total           6.8     93.1    564523  7687197
                                                        =============================================
                                                        okay.top n=1000; ave=2062; min,max=1333,31655; full=971; 
                                                        okay.all n=323341; ave=103; min,max=30,31655; full=267912; 
-----------------------------

Prior tr2aacds okayset classes (evg17pig3c/okayset/):
  Count   Class
  241182  alt
  323341  main+noclass
  --------------------
  564523  total

New trclass2pubset results, try9pig3c/publicset/pig3c.pubids
  Count   Class      Count   Cull-Class
  537569  alt        70344   cullalt
  241853  main           0   cullmain   # later will test mains for lowqual culls
   90224  noclass   303995   cullnoclass  
  ---------------------------------------
   869646 total     374339   cull-total

Totals for main+noclass putative gene loci are nearly same in prior and new,
update retains many more putative alternates.  The largest excess is for short-protein (<100aa)
models, which require either homology/external evidence to evaluate, or a subjective criterion
to ignore or cull as uninteresting.

2nd stage self-referential reclassification log
  altreclass: opts pDemoteAaPoor=0.9, pDemoteAntiSense=0.95, pDemoteSplitmap=0.8,
    pPromoteAaRef=0.5, pPromoteAaCons=0.1, pDemoteNoncode=0.8, mincov=75, altrenum=1, noclasscut=60
  readTrClass(pig3c.trclass)= 1243985 ** Same as tr2aacds okayset, pig3c.trclass.sum.txt
  readAaQual(.//inputset/pig3c.aa.qual)= 8429111, includes not coding-potential score
  readExontab(tmpfiles/pig3cnrcd1x.exontab)= 1243984, new alternate exon pattern data
  altreclass: nin/out=1243985/1243985, ndrop/keep=0/1243985, nrenum=419812, ngenediff=67897/636072
  -- not listed yet: ncull=374339

=cut

#====================================================================================
