## Supplementary material for "Longest protein, longest transcript or most expression, for accurate gene reconstruction of transcriptomes?": supOCC_stdprotein_calcs_compared.txt

Supplemental Document OCC. 
ORF computations comparison details (reference check codes, method command calls).
for Table AC8. Recovery of reference proteins by ORF computations from human and plant reference mRNA. 

There are several ways to compute proteins from mRNA transcripts, with differing results.
Two classes are (a) basic codon lookup scan of RNA sequence for longest open reading
frames (ORFs), and (b) prediction from statistical models of coding sequences, including
coding/non-coding metrics, intron/exon models, sequence signals. One should be aware of
what a protein translator is doing, in particular predictors miss, or mis-model from 20%
to 35% of standard reference proteins, while codon lookup calculations recover nearly all
reference proteins.

  Protein ORF computations for Human RefSeq and  Arabidopsis reference genes
  2019 January, Don Gilbert

                    Plant Ref        Human Ref
     ORF_calc     Equal  N.ORFs    Equal  N.ORfs    ORF_type
    -------------------------------------------------------------
    stdorf_ncbi    98%   48,359     97%   113,607   codon_lookup
    stdorf_evg     98%   48,349     97%   113,607   codon_lookup
    bestorf_evg    96%   48,349     93%   113,607   codon_lookup
    gmark_gmp      77%   47,892     69%   143,455   CDS_predict
    gmark_gmst     80%   45,830     71%   112,660   CDS_predict
    trandec_pred   83%   77,414     73%   113,607   CDS_predict
    trandec_long   87%  492,753     75% 2,735,910   codon_lookup
    -------------------------------------------------------------

This table shows the percentage of matches to reference proteins that ORF computations
will produce from the mRNA sequences of those same reference genes. The codon-lookup
translators have few mismatches (3% to 6%) to the reference, while the predictors have
large degrees of mis-match (20% to 35%) to reference.

For most uses with transcriptomes, the simpler codon lookup for longest ORF is the best
first step, and will match public gene/protein databases. Various coding/non-coding
metrics can be applied after basic ORF computations, and homology to reference genes are
measured, as part of gene set validations. In my experience the non-coding tests make
mistakes, calling known coding genes as non-coding often enough that you want to defer
this metric until after measuring for for known protein homology.

Reference proteins and mRNA sets (mRNA input to orf calcs)
PlantRef=Arabidopsis Araport11_genes.201606.aa
    arath16ap 48,359 transcripts/proteins of 27,655 gene loci
HumanRef=NCBI RefSeq  Homo_sapiens/reference/GCF_000001405.38_GRCh38.p12/, 2018
    human18nc 81051 non-redundant proteins in 113607 mrna transcripts (~22,000 loci)

Three methods (evg,ncbi,trin_long) are standard codon table lookup translation of nucleic triplets
  to amino acid, tabulating longest ORF, and all ORFs as a first step.
Three methods (gmark_gmp, gmark_gmst, trin_pred) are predictors based on  statistical models of 
  coding sequences, including coding/non-coding metrics, intron/exon models, 
  sequence signal statistical modeling for HMM  prediction of genes on chromosomes.
  
ORFs are calculated from the public  mRNA sequences for listed references, then compared
to reference proteins of same mRNA, for identity (end-to-end), identical substring (one is
contained in the other), different in some other way, or missing test ORF. 
Ambiguous code/noncode sequences exist, some are true biology ("NORF" is one term), 
some artifacts or ncRNA with computable ORF.
  
ORF calculation details: 
 stdorf_ncbi: NCBI  orf calc of ncbi toolkit 
  See https://www.ncbi.nlm.nih.gov/orffinder/
  used thru tbl2asn, "ncbibin/tbl2asn  -kc -Mt  -Vb -i human18nc.mrna"
 See also EMBOSS getorf, others.

 stdorf_evg: evigene/cdna_bestorf.pl -noutrorf -fullorf -cdna human18nc.mrna
  stdorf_evg choice is always complete protein, if it exists.  This matches well to reference
  gene sets, but will lose homolog of partial proteins from fragment mRNA assemblies.

 bestorf_evg: evigene/cdna_bestorf.pl -cdna human18nc.mrna   defaults are: -utrorf -bestorf
  bestorf_evg choice is between longest complete protein, or a longer partial protein,
  determined by ratio of sizes ( complete >= 85%  of longest is default).
 
 trandec_long: TransDecoder.LongOrfs : all ORFs/mRNA of good size ( ~10 ORF per transcript)
  this is a near equivalent to evg, ncbi basic ORF codon lookup, but returns most computed
  orfs per mRNA, rather than longest, and does not chose shorter full orfs over partials.

 trandec_pred: TransDecoder.Predict : uses statistical modeling of mRNA sequence, attempts to
  discriminate coding vs non-coding ORFs.
  TransDecoder is found at https://github.com/TransDecoder/TransDecoder/wiki
  
 gmark_gmp: genemark_suite/gmsn.pl --euk --faa human18nc.mrna
  GeneMark statistical modeling of coding-sequence genes on chromosomes, eukaryote variant.
 gmark_gmst: genemark/gmst for transcripts, gmst.pl --faa human18nc.mrna
  GeneMark modified for HMM-based modeling of coding-sequence on transcript sequences.
  Both from http://exon.gatech.edu/GeneMark/
  
See also http://eugenes.org/EvidentialGene/evigene/docs/evigene_vs_transdecoder_arabidopsis.txt
#-------------------------------------------------------

Code for ORF to Reference protein comparison statistics:

1.  Simple count of identical proteins for reference.aa x test.aa, orfcalcs/refequal.sh
usage:
atest=arath16ap_gmp;  aref=../Araport11_genes.201606;
env st=aa ref=$aref ./refequal.sh $atest  >> refequals.txt
See below refequal.sh

2.  Measure differences between reference.aa x test.aa, orfcalcs/aadiffu.pl
usage:
atest=arath16ap_gmp;  aref=../Araport11_genes.201606;
env idpre=AT dshow=0 ./aadiffu.pl $atest.aa $aref.aa > $atest.ref.aadiffu

aadiffu summary table (add to aadiffu.pl?):

perl -ne '($rd,$cla,$cw,$rw,$td)=@v=split; ($gn,$ti)=$rd=~m/^(\w+)\.(\d+)/;  map{ s/^[cr]://; }($cw,$rw); $gnok{$cla}{$gn}++; $gnokm{$cla}{$gn}++ if($ti==1); for $cc ($cla,"total") { $sum{$cc}{n}++; $sum{$cc}{rw}+=$rw; $sum{$cc}{cw}+=$cw; $sum{$cc}{dw} += $cw - $rw; $sum{$cc}{nalt}++ if($ti>1); } $nt++; END{ stats(); } sub stats { @k=qw(n nalt rw cw dw); @c=sort keys %sum; for $c (@c){ ($n,$nalt,$rw,$cw,$dw)=map{ $sum{$c}{$_}||0 }@k; $n||=1; ($pn,$pna)=map{ int(1000*$_/$nt)/10 }($n,$nalt); ($arw,$acw,,$adc)=map{ int(0.5+100*$_/$n)/100 } ($rw,$cw,$dw);  @gn=sort keys %{$gnok{$c}}; @gnm=sort keys %{$gnokm{$c}}; $ngn=@gn; $ngnm=@gnm;  print join("\t",$c,$n,$nalt,$dw,$pn,$pna,$arw,$acw,$adc,$ngn,$ngnm)."\n"; } } BEGIN{ print join("\t",qw(Class Nc Nalt dC_R pNc pNa aveRl aveCl avdCR nGn nG1))."\n"; } ' \
 $pt.ref.aadiff 

#-------------------------------------------------------
stdorf recheck, 19.oct.16, for Supplemental Document OCC.

recalc using refequal.sh, fastanrdb calc of identical proteins
Note diff for gmark, trandec w/ below aadiffu 
is due to nr-aa:  81000 nr-aa vs 113,000 all aa

Table SAC8a. Recovery of reference proteins by ORF computations from human RefSeq mRNA,
             counting unique, non-redundant proteins. I.e. 113607 RefSeq mRNA transcripts 
             contain 81051 unique proteins.
                       Human Ref (n=81051)
     ORF_calc       Equal% Equal.N   N.ORfs  
    ------------------------------------------
    stdorf_ncbi      96.9%  78,540   81,169 
    stdorf_evg       96.9%  78,554   81,174 
    bestorf_evg      93.4%  75,731   82,571 
    gmark_gmp        68.6%  55,573   97,247 
    gmark_gmst       71.4%  57,910   83,998 
    trandec_pred     72.6%  58,868   85,693 
    trandec_long     75.4%  61,095  748,481  
    ------------------------------------------

#-----------------------------------------------------
aacorftest/human18nc_evgorf_vs_other.info  

 orf computations vs Human RefSeq (2018 from NCBI) ref
 2019 January
 human ref data: Homo_sapiens/reference/GCF_000001405.38_GRCh38.p12/, 
    human18nc.aa human18nc.mrna (for orf compute)
    extracted from GenBank flatfile GCF_000001405.38_GRCh38.p12_rna.gbff.gz
 plant ref data: Araport11_genes.201606.aa, Araport11_genes.201606.cdna (for orf compute)
    arath16ap 48,359 transcripts/proteins of 27,655 gene loci

 evgorf = evigene cdna_bestorf.pl, with --noutrorf --fullorf (eof), or best full%part orf (eox) = tr2aacds
 ncbi   = NCBI standard orf calc, from tbl2asn ($nbin/tbl2asn  -kc -Mt  -Vb -i human18nc.mrna)
 gmp  = genemark_suite/gmsn.pl hmm gene predictor,  --euk --faa human18nc.mrna
 gmst = genemark/gmst for transcripts, gmst.pl -faa human18nc.mrna

 ref.aadiffu = classify ref x test for same (identical), calcsub, refsub, diff, miss
     calcsub = testorf substring of ref.aa, refsub = ref.aa substring of testorf

------- summary for human ref NCBI RefSeq 2018  ----
human18nc_ncbi  96 calcsub;     3191 diff;      11 miss;        500 refsub;     109809 same;
human18nc_evgf  96 calcsub;     3188 diff;      0 miss;         500 refsub;     109823 same;
human18nc_evgb  71 calcsub;     3775 diff;      0 miss;         4178 refsub;    105583 same;
human18nc_gmp   12931 calcsub;  11872 diff;     1666 miss;      16262 refsub;   70876 same;
human18nc_gmst  11660 calcsub;  2139 diff;      948 miss;       17242 refsub;   81618 same;
human18nc_trdec 59 calcsub;     4224 diff;      0? miss;        24129 refsub;   85195 same;
human18nc_trlorf 59 calcsub;    703 diff;       0 miss;         24127 refsub;   88718  same;
# added _trdec 2019.06 and trinlorf ? should add test orf count, e.g. trlorf has 20+ orfs per ref

------- summary for arabidopsis ref of Araport 2017 ----
arath16ap_ncbi  53 calcsub;     675 diff;       0 miss;         38 refsub;      47593 same;
arath16ap_evgf  50 calcsub;     668 diff;       10 miss;        38 refsub;      47593 same;
arath16ap_evgb  35 calcsub;     747 diff;       10 miss;        919 refsub;     46648 same;
arath16ap_gmp   4099 calcsub;   1449 diff;      2739 miss;      2915 refsub;    37157 same;
arath16ap_gmst  3092 calcsub;   396 diff;       2529 miss;      3408 refsub;    38934 same;
arath16ap_trdec  15 calcsub;     2367 diff;      1363 miss;      4580 refsub;    40034 same 
arath16ap_trlorf 22 calcsub;     114 diff;       25 miss;        6219 refsub;    41979 same; 

------- summary for mouse RefSeq mouse_GRCm38p4.rna 2016 ---
mouse16nc_evgb    91 calcsub; 1407 diff; 2326 refsub; 72379 same
mouse16nc_trdec   67 calcsub; 777 diff; 15340 refsub; 59782 same
mouse16nc_trlorf  63 calcsub; 314 diff; 15379 refsub; 60447 same  

Note: evgb vs evgf orf differs largely at refsub, due to existance of 5'orf leaders ahead of
true protein start, mis-estimated as partial5.  Should fix this in evigene bestorf.pl
Transdec may have same problem (higher refsub due to partial5 extention?)

#-----------------------------------------------------
#--- human detail stats from aadiffu ---

Class   Nc      Nalt    dC_R    pNc     pNa     aveRl   aveCl   avdCR   nGn     nG1
calcsub 12931   0       -528159 11.3    0       684.12  643.27  -40.83  1       0
diff    11872   0       -5608086        10.4    0       615.51  143.13  -472.37 1       0
miss    1666    0       -236522 1.4     0       141.97  0       -141.96 1       0
refsub  16262   0       715161  14.3    0       592.3   636.28  43.98   1       0
same    70876   0       0       62.3    0       701.77  701.77  0       1       0
total   113607  0       -5657606        100     0       666.87  617.07  -49.79  0       0
OF aacgmark/human18rna_gmp

Class   Nc      Nalt    dC_R    pNc     pNa     aveRl   aveCl   avdCR   nGn     nG1
calcsub 96      0       -7779   0       0       445.86  364.83  -81.02  1       0
diff    3191    0       124475  2.8     0       168.89  207.9   39.01   1       0
miss    11      0       -375559 0       0       34141.73        0       -34141.72       1       0
refsub  500     0       22058   0.4     0       471.32  515.43  44.12   1       0
same    109809  0       0       96.6    0       679.07  679.07  0       1       0
total   113607  0       -236805 100     0       666.87  664.78  -2.07   0       0
OF aacncbi/human18nc_gbf

Class   Nc      Nalt    dC_R    pNc     pNa     aveRl   aveCl   avdCR   nGn     nG1
calcsub 96      0       -7779   0       0       445.86  364.83  -81.02  1       0
diff    3188    0       157623  2.8     0       158.56  208     49.44   1       0
refsub  500     0       22058   0.4     0       471.32  515.43  44.12   1       0
same    109823  0       0       96.6    0       682.71  682.71  0       1       0
total   113607  0       171902  100     0       666.87  668.38  1.51    0       0
OF aacevg/human18nc_evgfo

Class   Nc      Nalt    dC_R    pNc     pNa     aveRl   aveCl   avdCR   nGn     nG1
calcsub 59      0       -5449   0       0       467.49  375.14  -92.35  1       0
diff    4224    0       234908  3.7     0       164.86  220.47  55.61   1       0
refsub  24129   0       995085  21.2    0       669.77  711.01  41.24   1       0
same    85195   0       0       74.9    0       691.07  691.07  0       1       0
total   113607  0       1224544 100     0       666.87  677.65  10.78   0       0
OF human18nc_transdec_pred

Class   Nc      Nalt    dC_R    pNc     pNa     aveRl   aveCl   avdCR   nGn     nG1
calcsub 59      0       -5449   0       0       467.49  375.14  -92.35  1       0
diff    703     0       44847   0.6     0       182.12  245.91  63.79   1       0
refsub  24127   0       994990  21.2    0       669.75  710.99  41.24   1       0
same    88718   0       0       78      0       670.06  670.06  0       1       0
total   113607  0       1034388 100     0       666.87  675.97  9.1     0       0
OF human18nc_transdec_all

#-----------------------------------------------------
#--- plant detail stats from aadiffu ---

SUM Stats for aacevgorf/arath16ap_efo
Class   Nc      Nalt    dC_R    pNc     pNa     aveRl   aveCl   avdCR   nGn     nG1
calcsub 50      7       -6855   0.1     0       515.84  378.74  -137.09 50      43
diff    668     238     16338   1.3     0.4     130.33  154.79  24.46   519     430
miss    10      0       -52     0       0       5.2     0       -5.19   10      10
refsub  38      31      737     0       0       448.87  468.26  19.39   38      7
same    47593   20464   0       98.4    42.3    435.48  435.48  0       27234   27129
total   48359   20740   10168   100     42.8    431.27  431.48  0.21    27655   27619
--------------------
SUM Stats for aacevgorf/arath16ap_eox
Class   Nc      Nalt    dC_R    pNc     pNa     aveRl   aveCl   avdCR   nGn     nG1
calcsub 35      3       -5613   0       0       640.77  480.4   -160.36 35      32
diff    747     260     31765   1.5     0.5     119.13  161.65  42.52   589     487
miss    10      0       -52     0       0       5.2     0       -5.19   10      10
refsub  919     355     41582   1.9     0.7     179.31  224.56  45.25   777     564
same    46648   20122   0       96.4    41.6    441.17  441.17  0       26787   26526
total   48359   20740   67682   100     42.8    431.27  432.67  1.4     27655   27619
--------------------
SUM Stats for aacgenemark/arath16ap_gmp
Class   Nc      Nalt    dC_R    pNc     pNa     aveRl   aveCl   avdCR   nGn     nG1
calcsub 4099    2292    -100255 8.4     4.7     463.67  439.21  -24.45  2804    1807
diff    1449    1036    -446723 2.9     2.1     476.29  167.99  -308.29 1060    413
miss    2739    602     -210751 5.6     1.2     76.94   0       -76.93  2266    2137
refsub  2915    1310    65882   6       2.7     364.75  387.35  22.6    2359    1605
same    37157   15500   0       76.8    32      457.28  457.28  0       22460   21657
total   48359   20740   -691847 100     42.8    431.27  416.96  -14.3   27655   27619
--------------------
SUM Stats for aacgmarkst/arath16ap_gmst
Class   Nc      Nalt    dC_R    pNc     pNa     aveRl   aveCl   avdCR   nGn     nG1
calcsub 3092    1807    -62602  6.3     3.7     470.58  450.34  -20.24  2108    1285
diff    396     155     8667    0.8     0.3     200.05  221.93  21.89   325     241
miss    2529    511     -187479 5.2     1       74.13   0       -74.12  2123    2018
refsub  3408    1568    95461   7       3.2     384.69  412.7   28.01   2686    1840
same    38934   16699   0       80.5    34.5    457.78  457.78  0       23026   22235
total   48359   20740   -145953 100     42.8    431.27  428.25  -3.01   27655   27619
--------------------
SUM Stats for aacncbix/arath16ap_gbf
Class   Nc      Nalt    dC_R    pNc     pNa     aveRl   aveCl   avdCR   nGn     nG1
calcsub 53      7       -6994   0.1     0       489.32  357.36  -131.95 53      46
diff    675     239     16160   1.3     0.4     127.36  151.3   23.94   525     436
refsub  38      31      737     0       0       448.87  468.26  19.39   38      7
same    47593   20463   0       98.4    42.3    435.5   435.5   0       27234   27130
total   48359   20740   9903    100     42.8    431.27  431.47  0.2     27655   27619
--------------------
SUM Stats for aactrindec/arath16ap_trindec
Class   Nc      Nalt    dC_R    pNc     pNa     aveRl   aveCl   avdCR   nGn     nG1
calcsub 15      0       -4386   0       0       571.67  279.27  -292.39 15      15
diff    2367    989     -581189 4.8     2       313.26  67.73   -245.53 1959    1378
miss    1363    195     -142043 2.8     0.4     104.21  0       -104.2  1250    1168
refsub  4580    1903    112662  9.4     3.9     401.28  425.88  24.6    3697    2677
same    40034   17653   0       82.7    36.5    452.76  452.76  0       23278   22381
total   48359   20740   -614956 100     42.8    431.27  418.55  -12.71  27655   27619
--------------------
SUM Stats for aactrindec/arath16ap_trinlorf
Class   Nc      Nalt    dC_R    pNc     pNa     aveRl   aveCl   avdCR   nGn     nG1
calcsub 22      0       -5144   0       0       669.23  435.41  -233.81 22      22
diff    114     21      1423    0.2     0       193.51  205.99  12.48   109     93
miss    25      0       -330    0       0       13.2    0       -13.19  25      25
refsub  6219    2686    153211  12.8    5.5     398.86  423.49  24.64   4733    3533
same    41979   18033   0       86.8    37.2    436.84  436.84  0       24841   23946
total   48359   20740   149160  100     42.8    431.27  434.35  3.08    27655   27619
--------------------

#-------------------------------------------------------
#!/bin/bash
## refequal.sh
## defaults: cds seqtype, ref for arabid, human

# check if fastanrdb on path
fnrdbok=`which fastanrdb`
if [ $? != 0 ]; then
  xnrpath="$HOME/bio/apps/exonerate/bin";
  if [  -d $xnrpath -a -x $xnrpath/fastanrdb ]; then
    export PATH=$xnrpath:$PATH
  fi
fi

tes=$1
if [ "X$st" = "X" ]; then st=cds; fi
if [ "X$ref" = "X" ]; then echo "#ERR: missing env ref=file"; exit -1; fi
if [ "X$reft" = "X" ]; then reft=ref_; fi

if [ -f $reft$tes.nr$st.ids ]; then
  echo "# have $reft$tes.nr$st.ids "
elif [ -f $tes.$st ]; then
  if [ ! -f $ref.$st ]; then echo missing $ref.$st; exit -1; fi
  cat $ref.$st | sed 's/^>/>REF/;' > $reft$tes.aatwo
  cat $tes.$st | sed 's/^>/>TEST/; s/\*$//; ' >> $reft$tes.aatwo
  fastanrdb $reft$tes.aatwo | grep '>' > $reft$tes.nr$st.ids
  /bin/rm $reft$tes.aatwo
else
  echo missing $tes.$st
  echo usage: env st=cds ref=refset/ref_arath16apnr ./refequal.sh bestevg_arath7in
  echo counts identical sequences of bestevg_arath7in.cds to ref_arath16apnr.cds
fi

if [ -f $reft$tes.nr$st.ids ]; then
  nref=`grep -c REF $reft$tes.nr$st.ids`
  ntest=`grep -c TEST $reft$tes.nr$st.ids`
  neq=`grep REF $reft$tes.nr$st.ids | grep TEST | wc -l`
  pref=`echo "scale=2; 100*$neq/$nref;" | bc`
  echo  "equal_ref$tes   neq=$neq, pref=$pref%, nref=$nref, ntest=$ntest"
fi

#-------------------------------------------------------
